# Space-time resolved inference-based whole-brain neurophysiological mechanism imaging: application to resting-state alpha rhythm

**DOI:** 10.1101/2022.05.03.490402

**Authors:** Yun Zhao, Mario Boley, Andria Pelentritou, Philippa J. Karoly, Dean R. Freestone, Yueyang Liu, Suresh Muthukumaraswamy, William Woods, David Liley, Levin Kuhlmann

## Abstract

Neural mechanisms are complex and difficult to image. This paper presents a new space-time resolved whole-brain imaging framework, called Neurophysiological Mechanism Imaging (NMI), that identifies neurophysiological mechanisms within cerebral cortex at the macroscopic scale. By fitting neural mass models to electromagnetic source imaging data using a novel nonlinear inference method, population averaged membrane potentials and synaptic connection strengths are efficiently and accurately imaged across the whole brain at a resolution afforded by source imaging. The efficiency of the framework enables return of the augmented source imaging results overnight using high performance computing. This suggests it can be used as a practical and novel imaging tool. To demonstrate the framework, it has been applied to resting-state magnetoencephalographic source estimates. The results suggest that endogenous inputs to cingulate, occipital, and inferior frontal cortex are essential modulators of resting-state alpha power. Moreover, endogenous input and inhibitory and excitatory neural populations play varied roles in mediating alpha power in different resting-state sub-networks. The framework can be applied to arbitrary neural mass models and has broad applicability to image neural mechanisms in different brain states.

**Highlights:**

- The whole-brain imaging framework can disclose the neurophysiological substrates of complicated brain functions in a spatiotemporal manner.
- Developed a semi-analytical Kalman filter to estimate neurophysiological variables in the nonlinear neural mass model efficiently and accurately from large-scale electromagnetic time-series.
- The semi-analytical Kalman filter is 7.5 times faster and 5% more accurate in estimating model parameters than the unscented Kalman filter.
- Provided several group-level statistical observations based on neurophysiological variables and visualised them in a whole-brain manner to show different perspectives of neurophysiological mechanisms.
- Applied the framework to study resting-state alpha oscillation and found novel relationships between local neurophysiological variables in specific brain regions and alpha power.

## Introduction

A long-standing goal in neuroscience is to image the human brain’s neural mechanisms to understand the patterns of neural activity that give rise to mental processes and behaviours. Modern human neuroimaging methods have made significant progress towards unlocking these mechanisms. Functional magnetic resonance imaging (fMRI), the most commonly used functional neuroimaging technique, has revealed fundamental insights into spontaneous and task-dependent brain activity by imaging the blood oxygenation changes related to the metabolic processes of neurons^1^. Electroencephalography (EEG) and magnetoencephalography (MEG) have been used to study oscillatory neural activity by measuring the electrical and magnetic fields induced by source currents within the brain^2^. However, both fMRI and M/EEG do not provide a direct measure of the neurophysiological variables that generate these source currents or blood oxygenation changes^3, 4^. In particular, it is well known that cellular membrane potentials and post-synaptic potentials are critical neurophysiological variables involved in dynamic information processing in the brain^5^ and the generation of macroscopic currents^2^ and modulation of blood-oxygenation signals^1^. These neurophysiological variables have long been the purview of cellular and systems neuroscience and have afforded the elucidation of the neural mechanisms of many brain functions^5^. While there have been advances with multi-photon imaging of neural tissue^6^ and magnetic resonance spectroscopy to image specific neurotransmitter changes^7^, work remains to be completed to provide efficient space-time resolved imaging of the aforementioned neurophysiological variables across the whole human brain.

Computational models provide an alternative to capture neural mechanisms by abstracting latent population averaged neurophysiological variables in mathematical models^8^. Neural mass models (NMMs) are neural population-level models used in studying neural oscillations by modelling the activity of groups of excitatory and inhibitory neurons within a brain region and approximating the averaged properties and interactions^8^. The combination of such models and model inversion, which estimates neurophysiological variables in the model from data^9^, provides a powerful way to infer neural mechanisms from functional neuroimaging recordings. Dynamic causal modelling (DCM) has been one of the principal frameworks that study neural mechanisms in this way^10, 11^. Specifically, DCM focuses on the task-dependent effective connectivity among multiple selected brain regions. Given the challenge of model inversion for highly complex systems like the brain, there has been little work done to image space-time resolved brain-wide neurophysiological changes. Model inversion that can scale to the whole brain at a spatial resolution commensurate with current human neuroimaging techniques, is a computationally formidable challenge for complex non-linear neural models^9^. Most model inversion methods that are efficient and therefore scalable, usually rely on linearising nonlinear neural models and are therefore less accurate^12, 13^. On the other hand, inversion methods that are accurate, like sampling techniques, are usually computationally demanding and therefore less scalable^14, 15^. Thus, existing methods are not suitable for space-time resolved whole-brain imaging with a reasonable computational time budget and acceptable estimation accuracy.

This paper presents a novel whole-brain imaging framework, Neurophysiological Mechanism Imaging (NMI), that extends existing electromagnetic source imaging methods^16^ to infer and image neurophysiological variables in a space-time resolved manner. The framework is demonstrated by applying it to human MEG resting-state alpha (8-12 Hz) rhythm data and inferring and imaging population mean (i.e., spatial, not time averaged) membrane potentials and population mean synaptic strengths of cerebral-cortical excitatory and inhibitory neuronal populations at the spatial and temporal resolutions afforded by MEG (6mm spaced sources, 400 Hz sampling rate). The framework allows for the general application of NMMs containing an arbitrary combination of interconnected excitatory and inhibitory populations and involves an efficient time-domain Bayesian inference method derived specifically for nonlinear NMMs.

Although the accuracy and interpretation of the inferred neurophysiological variables of such a framework depends on the assumed neural model applied in a specific situation^9^, a similar problem is present in standard fMRI analysis where a hemodynamic response function (HRF) is applied even though it is known to depend on various factors and therefore can be modelled in different ways^17^. The typical solution to get around these complexities is to apply a canonical HRF to each voxel in the brain, and this has enabled the general application of fMRI to many different neuroscience topics. With this in mind, and akin to DCM for EEG/MEG^11^, here we present a novel imaging framework using a canonical NMM. Moreover, although the framework can theoretically work for large scale fully interconnected whole-brain models and infer any population averaged connection strength between any two neural populations, this paper assumes that each MEG-inferred source point in the brain is represented by a canonical NMM, whose linearly summed long-range inputs from the NMMs of other source points are treated as a single input to each canonical model. This helps circumvent the complexity of the solution space of inferring the fully interconnected whole-brain model, and provides an original space-time resolved view of neurophysiological variables in the human brain. In addition, through application of this novel method we consider different ways in which the inferred variables can be imaged and reveal new insights into the resting-state brain and alpha rhythm^18, 19^.

## Material and Methods

### Framework for inference-based whole-brain neural mechanism imaging

The proposed framework for quantifying and imaging neurophysiological variables, employs a canonical NMM^20^, an efficient Bayesian inference method, and multiple statistical analysis schemes that can be applied to any source time-series derived from electromagnetic imaging. Our method involves a fast, semi-analytic solution to handle the propagation of estimates through the non-linear neural model by re-deriving the well-known Bayesian inference method, Kalman filtering using the specific nonlinearities of the model without having to linearise the model around its equilibrium points^21^. This method is referred to here as the semi-analytic Kalman filter (AKF) and creates a more accurate and computationally efficient method which allows, in the examples presented here, the modelling of the whole brain with 4714 NMMs, and provides high spatial and temporal resolution imaging (6mm source spacing, 400 Hz sampling rate). To demonstrate the utility of the framework, we applied it to beamformed MEG recordings of healthy human male subjects^22^ to investigate the brain-wide neurophysiological mechanisms of eyes-closed resting-state alpha oscillations. Each MEG source time-series derived from beamforming is treated as proportional to the local extracellular dipole current formed by the combination of population averaged excitatory and inhibitory synaptic currents associated with cortical pyramidal cells^2, 23^. This combination of synaptic currents is treated as the output of a single NMM, such that one NMM is used per MEG source point to capture its underlying neural dynamics, as shown in **Fig. 1a**. By effectively inverting the model, estimates of neurophysiological variables can be obtained.

**Figure 1.**
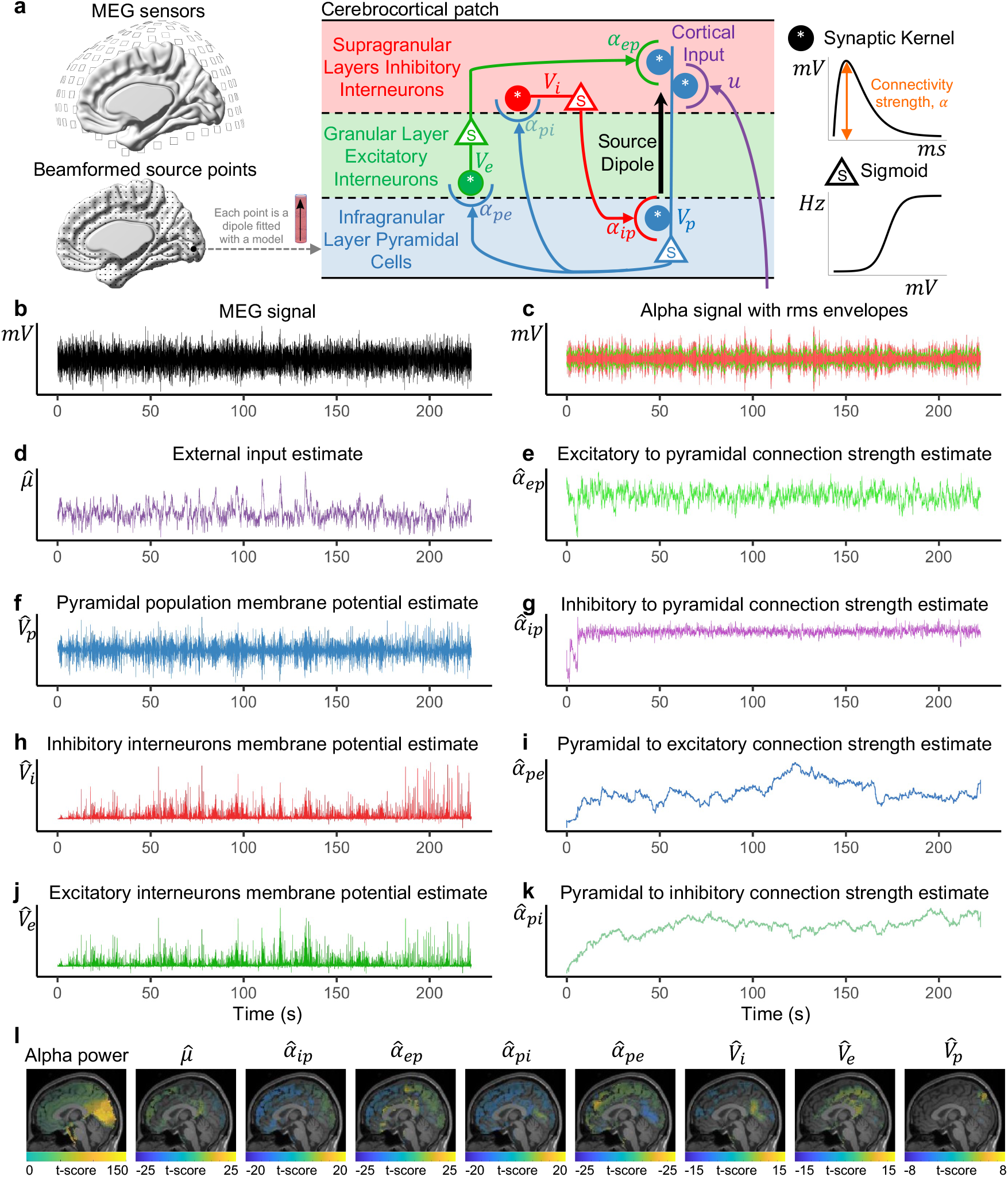
Schematic and example of space-time resolved inference-based whole brain imaging of neurophysiological mechanisms. (**a**) Schematic of whole brain modelling with the canonical NMM. Each MEG source point was fitted with an NMM. The canonical NMM used in this study contains three populations: pyramidal population, excitatory interneurons, and inhibitory interneurons. Each population is characterised by the mean membrane potentials *V*_*n*_, and each connection strength between two populations is represented by *α*_*mn*_. They were inferred from data by the AKF. (**b**) A sample recording from a MEG source point located in the occipital lobe of a single subject. (**c**) Alpha (8-12 Hz) band time-series in red extracted from the sample recording. The corresponding green lines, the rms envelopes, quantify the evolution of the signal power at a slower timescale. (**d**) – (**k**) The estimation timeseries of modelled neurophysiological variables obtained by feeding the sample recording into the AKF. (**l**) An example whole-brain contrast imaging shows the mean difference of alpha power (rms envelope) and neurophysiological variable estimates between two conditions: occipital strong and weak alpha oscillations. For each source point, a two-sample t-test was performed to evaluate the significance of the mean difference of the neurophysiological variable estimates between the aforementioned two conditions. The activated areas in the sub-images represent statistically significant difference was observed after corrections for multiple comparisons (significance level *α* = 0.05). See methods for more details about the statistical analysis scheme.

The NMM considered in this framework is illustrated in **Fig. 1a**. The model has three neural populations: excitatory pyramidal cells (p), spiny stellate excitatory cells (e) and inhibitory interneurons (i), which coarsely imitates the structure of a small patch of cerebral cortex^1-3^. The pyramidal population is driven by cortical input, *µ*, and excites the excitatory and inhibitory interneurons, while the excitatory and inhibitory interneurons provide excitatory and inhibitory feedback, respectively, to the pyramidal population. The cortical input encodes the neighbouring and distant afferent input to the cortical patch. The state of a neural population is denoted by the population mean membrane potential, *V*_*n*_. The population averaged connection strength between two neural populations is defined as *α*_*mn*_ (presynaptic and postsynaptic neural populations are indexed by *m* and *n*), representing lumped connectivities that incorporate average synaptic gain, number of connections, and averaged maximum firing rate of the presynaptic populations. The variables *V*_*n*_, *α*_*mn*_ and *µ* are the ones being estimated and imaged across the whole brain.

To estimate the neurophysiological variables of the NMMs, the AKF was applied to each of an individual’s 4714 source time-series obtained from resting eyes-closed MEG data. Below the data and methods used to illustrate inference-based whole-brain neural mechanism imaging at the individual and group level are described.

### Study population

The data used herein was collected from a study analysing resting state and anaesthetic-induced power changes in EEG/MEG signals. Here we summarise the key details of the data. For further information see previous papers^22, 50^. Twenty-two volunteers were recruited. All participants signed a written informed consent form prior to participation under the approved protocol (Alfred Hospital HREC approval number 260/12). Participants were right-handed adult males, aged between 20 and 40 years (average age was 24 years), and had a body mass index between 18 and 30. Contraindications to magnetic resonance imaging or magnetoencephalography (such as implanted metal foreign bodies) were an exclusion criterion. Candidates with neurological diseases, mental illnesses, epilepsy, heart disease, respiratory diseases, obstructive sleep apnea, asthma, motion sickness, and claustrophobia have been excluded from the study. In addition, any recent use of psychoactive drugs or other prescription drugs and any recreational drugs resulted in exclusion.

### MEG collection

Empirical eyes closed resting-state MEG data collection was carried out in a room shielded from magnetic or electrical interference (Euroshield Ltd., Finland). A 306-channel Elekta Neuromag TRIUX magnetoencephalography system (Elekta Oy, Finland) was used to record brain magnetic field activity at a sampling rate of 1,000 Hz. The system consists of 204 planar gradiometers and 102 magnetometers. Head position in relation to the recording system was recorded using five head-position indicator coils and was continuously monitored by measuring the magnetic fields produced by the coils in relation to the cardinal points on the head (nasion, and left and right preauricular points) which were determined before commencement of the experiment using an Isotrack 3D digitiser (Polhemus, USA). The internal active shielding system for 3D noise cancellation is disabled for subsequent source space analysis. Exactly five minutes of awake, eyes closed resting-state was collected for each participant.

### Structural T1-magnetic resonance imaging

To correlate brain function with structure, a single structural T1-weighted MRI scan was performed for each participant using a 3.0 TIM Trio MRI system (Siemens AB, Germany) with markers (vitamin E capsules) used to highlight the digitised fiducial points for the nasal apex and left and right preauricular points for subsequent source space reconstruction. T1-weighted images were acquired on a sagittal plane with a magnetisation prepared rapid gradient echo pulse sequence with an inversion recovery (176 slices: slice thickness, 1.0 mm; voxel resolution, 1.0 mm^3^; pulse repetition time, 1,900 ms; echo time, 2.52 ms; inversion time, 900ms; bandwidth, 170 Hz/Px; flip angle, 9°; field of view; 350 mm × 263 mm × 350 mm; orientation, sagittal; acquisition time, approximately 5 min).

### Pre-processing and source reconstruction

Data analysis was performed in Fieldtrip^51^ version 20200828 and custom MATLAB (MathWorks, USA) scripts and toolboxes. Magnetoencephalography data from 22 subjects was visually inspected to exclude any malfunctioning channels, and any artifactual segments resulting from eye movements or blinks, jaw clenches or movements, head movements, breathing, and other muscle artefacts were excluded. Magnetoencephalography data was filtered using the temporal signal-space separation algorithm of MaxFilter software version 2.2 (Elekta Neuromag Oy, Finland) and the signals from magnetometers and planar gradiometers were combined using Fieldtrip. All data was bandpass filtered at 1 to 100 Hz and any line noise at 50, 100, and 150 Hz was removed using notch filters. All recordings were visually inspected and any artefactual segments resulting from eye movements or blinks, jaw clenches or movements, head movements, breathing, and other muscle artefacts were excluded from further analysis. According to the minimum record length after data pre-processing, the MEG recordings of all subjects were trimmed to 3.75 minutes to limit bias to a particular subject in group analysis.

Co-registration of each subject’s magnetic resonance imaging to their scalp surface was performed using fiducial realignment, and boundary element method volume conduction models were computed using a single shell for MEG^52, 53^. MEG was spatially normalized to the MNI MRI template using the Segment toolbox in SPM8 to serve as a volume conduction model in subsequent source level analyses^54^. An atlas-based version of the linearly constrained minimum variance spatial filtering beamformer was used to project sensor level changes onto sources^16, 55^. Global covariance matrices for each band (after artifact removal) were derived, which were utilised to compute a set of beamformer weights for all brain voxels at 6-mm isotropic voxel resolution. Following inverse transformation, the 5061 voxels were assigned 1 of 90 automated anatomical labelling atlas labels (78 cortical, 12 subcortical) in the subjects’ normalised coregistered magnetic resonance image (based on proximity in Euclidean distance to centroids for each region) to identify regional source changes^56^. The resulting beamformer weights were normalised to compensate for the inherent depth bias of the method used^55^. Given that MEG is primarily sensitive to superficial brain structures it was decided to focus only on analysing the 4714 voxels in the grey matter of cerebral cortex. The beamformed MEG source time-series at each voxel was estimated in the direction of the most power and downsampled to 400 Hz to make neurophysiological variable inference tractable within a reasonable time budget and data storage, while maintaining a relatively high temporal resolution.

### Whole-brain imaging of neurophysiological mechanisms

As described above, at a high level the whole-brain imaging framework aims at efficiently imaging inferred population averaged neurophysiological variables across space and time at the resolution afforded by EEG or MEG inverse source reconstruction methods. In this paper, this is specifically achieved by augmenting MEG inverse source reconstruction methods by inferring neurophysiological variables from derived cortical source activity.

In the presented implementation, the proposed framework utilises a canonical NMM^20^ to approximate a small cortical patch which contains multiple neuronal populations characterised by population averaged mean membrane potentials, and the interplay between populations quantified by the population averaged synaptic connectivity strengths. While the framework presented here is general enough to use arbitrarily complex NMMs. There are 4714 reconstructed MEG source points in this study and each point was fitted with an NMM. The local neural activity and inter-regional connections can be captured by each NMM and quantitatively reflected by the model’s neurophysiological variables. A novel data-driven Bayesian estimation scheme, namely the AKF, was employed to infer the model’s neurophysiological variables from the actual MEG-derived source time-series. Neural mechanisms were then characterised by calculating different statistics based on the neurophysiological variable estimates (details below). Whole-brain imaging was then achieved by projecting the statistics to the corresponding source point location on the MRI and interpolating the values to the neighbouring area. Overall, the proposed framework is a general protocol for working with different modalities of data such as MEG, EEG, Electrocorticography (ECoG), and supporting various estimation schemes such as unscented Kalman filter, particle filter, and imaging a variety of statistics to show different aspects of the physiological bases of the neural activity, and most importantly it can show the reasonably fine-grained spatial and very high temporal patterns of neural activity.

In this study, the whole-brain imaging framework has been applied to study the neurophysiological mechanisms underlying the resting-state alpha oscillation. Two conditions, contrast imaging and Pearson’s correlation-based imaging, were employed to illustrate the neurophysiological mechanisms from two statistical perspectives. To overcome the multiple comparisons problem, a nonparametric permutation test was used to determine the group-level significant voxels across the whole brain for each type of statistical analysis. By projecting the statistics of interest onto the MRI, one can observe the structure-function relationships associated with the inferred neurophysiological variables. Further details on the empirical data analysis are provided after first considering the canonical NMM employed, the mathematics underlying the AKF inference method, and simulations and analysis that verify the theoretical estimation accuracy and computational efficiency of the AKF.

### Neural mass model

It is rational to employ NMMs in the framework to model the macroscopic neural activity measured by M/EEG because the NMM is defined in line with the dynamics of cerebral cortex and the output of the model physiologically corresponds to M/EEG derived source-estimates^2^. EEG and MEG measure electric and magnetic fields, respectively, associated with postsynaptic potentials generated at the synaptic connections of large numbers of parallel pyramidal neurons whose activity is coherent in time and space. When presynaptic cells release excitatory neurotransmitters, such as glutamate, that bind to their associated post-synaptic receptors, they produce excitatory postsynaptic potentials (EPSPs) that act as current sinks. Conversely, inhibitory postsynaptic potentials (IPSPs) act as current sources and are generated by inhibitory neurotransmitters, such as gamma-aminobutyric acid (GABA), binding to their associated postsynaptic receptors that hyperpolarize postsynaptic cells. The summation of these sink and source currents generated by numerous parallel pyramidal cells gives rise to, as a first order approximation, the macroscopic source current dipoles inferred by electromagnetic source imaging^57^, In the current paper, source imaging has been derived by MEG beamforming as described above. The dynamics of the NMM are mathematically defined by differential equations that are dependent on neurophysiological variables.

The mathematical formulation given below is general enough to define any large scale NMM that consists of any arbitrary combination of interconnected excitatory and inhibitory neural mass populations. However, the implementation of the local canonical NMM used in this study is derived from the model introduced by Jansen and Rit^20^ and has been outlined in previous works^14, 21, 58^. The NMM is suitable to model MEG measured at this scale (virtual source dipoles with 6 mm spacing), in line with similar NMMs used to describe EEG or MEG^46, 59, 60^. A single independent NMM was fitted to each MEG source point (4714 models in total). NMMs were not coupled between source points and thereby, neurophysiological variables principally captured the local neural activity within the vicinity of a single cortical source point. The input parameter in **Fig. 1a**, *µ*, described neighbouring and distant afferent input to the patch approximated by a single NMM.

The Jansen and Rit model is composed of three neural populations (excitatory, inhibitory, and pyramidal). The pyramidal population (in infragranular layers) excites the spiny stellate excitatory population (in granular layer IV) and inhibitory interneurons (in supragranular layers), is driven by endogenous input, is excited by the spiny stellate excitatory population and inhibited by the inhibitory interneurons. Neural populations are described by their time varying mean (spatial, not time averaged) membrane potential, *V*_*n*_, which is the sum of contributing population mean post-synaptic potentials, *V*_*mn*_ (pre-synaptic and post-synaptic neural populations are indexed by *m* and *n*) and connected via synapses in which the parameter, *α*_*mn*_ quantifies the population averaged connection strength.

In general, a NMM is a time-varying dynamical system and each mean post-synaptic potential, V_*mn*_, is defined as two coupled, first-order, ordinary differential equations,

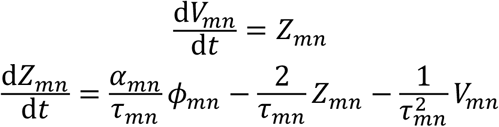

where *τ*_*mn*_ is a lumped time constant (population averaged synaptic response time constant) and *α*_*mn*_ is a lumped connection strength parameter that incorporates the average synaptic gain, the number of connections and the average maximum firing rate of the presynaptic populations. Both time constants and connection strengths are dependent on the type of presynaptic population. For example, GABAergic inhibitory interneurons typically induce a higher amplitude post-synaptic potential with a longer time constant than glutamatergic excitatory cells. *ϕ*_*mn*_ encodes the inputs to the population and it may come from external regions, *µ*, or from other intra-regional populations within the model, *g*(*V*_*m*_), where

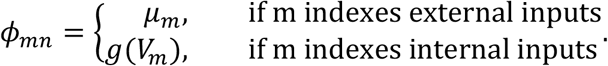

The various populations within the model are linked via the activation function, *g*(*V*_*m*_), that describes a mean firing rate as a function of the pre-synaptic population’s mean membrane potential. The activation function exploits a sigmoidal relationship (limited firing rate due to refractory period of the neurons) between the mean membrane potential and firing rate of each of the populations. This sigmoidal nonlinearity may take different forms, but for this study the error function form is used (as it facilitates the derivation of the AKF below) where

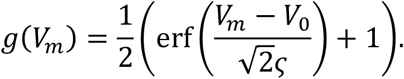

The quantity ς describes the slope of the sigmoid or, equivalently, the variance of firing thresholds of the presynaptic population (assuming a Gaussian distribution of firing thresholds). The mean firing threshold relative to the mean resting membrane potential is denoted by *V*_0_. The parameters of the sigmoidal activation functions, ς and *V*_0_, are usually assumed to be constants.

Depending on the inputs to a given population, the population averaged membrane potentials can be determined by the formula, *V*_*n*_ = ∑_*m*_*V*_*mn*_. Moreover, the postsynaptic potential, *V*_*mn*_ defined in the ordinary differential equations can conveniently be written as the convolution of the input firing rate, *ϕ*_*mn*_, with the postsynaptic response kernel, *h*_*mn*_,

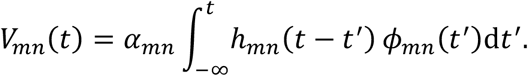

The post-synaptic response kernels denoted by *h*_*mn*_(*t*) describe the profile of the post-synaptic membrane potential of population *n* that is induced by an infinitesimally short pulse from the inputs (like an action potential). The post-synaptic response kernels are parameterised by the time constant *τ*_*mn*_ and are given by

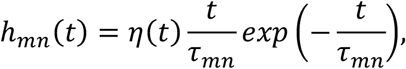

where *η*(*t*) is the Heaviside step function.

In summary, a single NMM component maps from a mean pre-synaptic firing rate to a post-synaptic potential. The terms of interest in our study are population mean membrane potentials, V_*n*_, and connectivity strengths, *α*_*mn*_, and cortical input, *µ*, which need to be inferred from the data while fixing the other parameters defined by Jansen & Rit^20^ and Freestone et. al^35^.

Below the NMM is expressed in matrix vector form to facilitate the exposition of the AKF in the following section and to highlight the generality of the framework. The general NMM is linked to the canonical NMM where appropriate. The state vector representing the postsynaptic membrane potentials is defined as

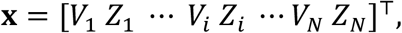

where the subscript, *i*, indexes the possible connections between populations in the neural mass. There are two states per population averaged synapse where V_*i*_ denotes the mean post-synaptic membrane potential and Z_*i*_ denotes the first order derivative of the mean post-synaptic membrane potential. Note that V_*i*_ can be interpreted as the vectorisation of V_*mn*_ and is distinct from V_*n*_. In the context of the implementation of each local canonical NMM, the state vector can be expressed as

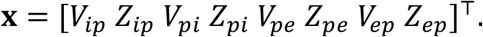

The general fully interconnected NMM can be expressed in a matrix notation

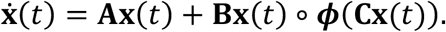

The matrix A encodes the dynamics induced by the membrane time constants. For *N* synapses, A has the block diagonal structure

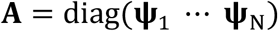

where

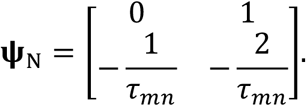

The matrix B maps the connectivity gains to the relevant sigmoidal activation function and is of the form

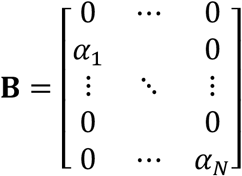

where α_*i*_ are the connection strength parameters in the NMM and can be thought of as the vectorisation of *α*_*mn*_. The vector function ***ϕ***() has the following form

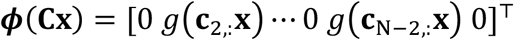

where *g*() is the sigmoid function defined above. The adjacency matrix, ***C***, defines the connectivity structure of the model. It is a matrix of zeros and ones that specifies all the connections between neural populations that has the block structure

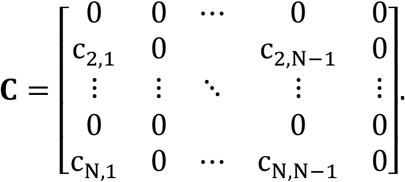

In general, for the AKF derivation below to work only connectivity strengths α_*i*_ can be estimated while the other parameters are treated as known constants. The benefit of this approach is that it leads to a highly efficient and scalable estimation algorithm. The connectivity strengths to be estimated can be formed into a parameter vector

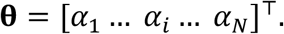

In the specific context of implementing the local canonical NMM that only involves the pyramidal, excitatory interneuron and inhibitory interneuron populations, the parameter vector for each canonical NMM can be expressed as.

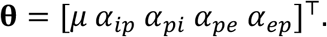

The dynamics for the parameters are modelled as constant

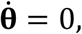

however, during estimation they are assumed to follow a random walk. The differential form of the parameter vector facilitates augmenting the parameters to the state vector for estimation purposes. The augmented state space vector is created as

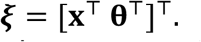

Accordingly, the matrices **A**_θ_, **B**_θ_, and **C**_θ_ are created based on **A**, **B**, and **C** to match the augmented state vector.

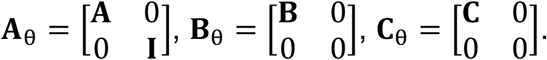

The augmented state space model is of the form

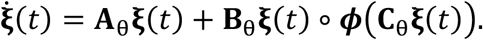

It is necessary to discretise the model with respect to time for estimation purposes. Therefore, rather than define the matrices **A**_θ_, **B**_θ_, and **C**_θ_ for continuous time, the discrete time formulation is given. The Euler method was used for discretising the model. For the AKF it is also necessary to model uncertainty in the model by an additive noise term. By including a Gaussian white noise term with zero mean and known covariance matrix **Q**, the discrete time augmented state space model is denoted by

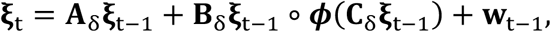

where ξ ∈ ℝ^N×1^ and the discrete time version matrices **A**_δ_, **B**_δ_, **C**_δ_ are ∈ ℝ^N×N^ and have the form

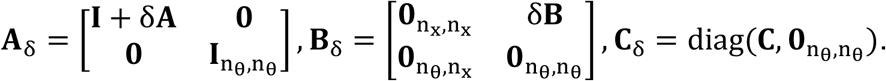

In this study, N = 13 is the total number of elements in the augmented state vector, *n*_x_ = 8 is the number of model states, and *n*_θ_ = 5 is the number of model parameters.

In forward models, **w**_t −1_ can be used as a driving term to simulate unknown input to the system from afferent connections or from other cortical regions. However, for model inversion purposes, this additional term also facilitates estimation and tracking of parameters via Kalman filtering or other Bayesian inference schemes. For the Kalman filter, the covariance of **w**_t −1_ quantifies the error in the predictions through the model. If one believed the model is accurate, then one would set all the elements of **Q** to a small value. On the other hand, a high degree of model-to-brain mismatch can be quantified by setting the elements of **Q** to larger values. According to previous works^14, 21^, a small constant value, 5µV, was used for the model uncertainty, which prevents the filter converging, and enabling new measurements to continue to influence the estimation.

### Model of MEG/EEG measurements

In general, M/EEG-derived source time-series for each source point can be treated as proportional to the combination of the mean excitatory and inhibitory postsynaptic contributions to the local pyramidal population’s mean membrane potential. The measurement model that relates the M/EEG-derived source time-series to the augmented state vector, ξ_t_, is given by

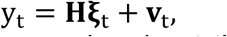

where y_t_ is the vector of M/EEG source signals at time *t*, and v_t_∼𝒩(0, **R**) is a zero mean, spatially and temporally white Gaussian noise process with a standard deviation of 1 mV^35^, that simulates measurement errors. For model inversion purposes, the variance of v_*t*_ quantifies the confidence we have in the measurements. The matrix **H** when multiplied with the augmented state vector defines a summation of the post-synaptic membrane potentials (corresponding to pyramidal populations) that contribute to each M/EEG source point. Within the context of applying the framework to the canonical NMM, the formulation for each local NMM can be simplified such that y_t_ can be considered a univariate source time-series for each independent NMM and **H** reduces to a vector. This results in 4714 decoupled NMMs where the local connections are still present and external connections are treated as a lumped input, and 4714 separate estimators. For example, the measurement model of a single NMM (excluding the noise term) is defined as

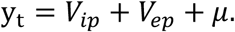

### Neurophysiological variable estimation with analytical Kalman Filter (AKF)

In general, Kalman filters provide the minimum mean squared error estimates for the augmented state vector, under the assumption that they are normally distributed. Assumptions about normality are reasonable if considering the central limit theorem and the population level neurophysiological variables as a mean of the many underlying microscopic neurophysiological variables. As illustrated in **Supplementary Fig. 5** the prediction errors of the AKF when applied to real data follow close to a normal distribution, suggesting assumptions about normality are valid. Moreover, without the normality assumption it would be difficult to obtain such an efficient estimation method.

The AKF as presented here is a highly stable and accurate iteration on prior works^21, 35^ that evolved from attempting to follow the same derivation of the original Kalman filter for linear systems, but instead deriving the filter for general nonlinear NMMs using the specific sigmoidal non-linearity given above. Thus, the AKF can be thought of as an assumed density filter. Mathematically stated, the aim of the estimation is to calculate the most likely posterior distribution of the augmented state given the previous measurements,

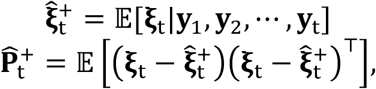

which are known as the a posteriori state estimate and state estimate covariance, respectively. In prior work attempting to derive the AKF some of the terms of the state and covariance estimate calculations were difficult to derive analytically and so had to be calculated using the unscented transform as used in the unscented Kalman filter (UKF) for nonlinear systems^41^. However, in the current instantiation of the AKF these mathematical difficulties have been overcome and all terms except one are calculated analytically. The remaining term is calculated semi-analytically using the Error Function. Moreover, the algorithm has been made more numerically stable by including iterative optimisation techniques^61^ to ensure the estimated covariance matrix remains positive semi-definite. This leads to a highly stable, accurate and computationally efficient time-domain estimation method.

The estimator proceeds in two stages: prediction and update. In prediction, the prior distribution (obtained from the previous estimate) is propagated through the neural mass equations. This step provides the so called *a priori* estimate, which is a Gaussian distribution with mean and covariance,

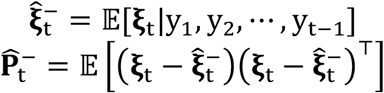

In the second stage, the a posteriori state estimate is calculated by correcting the a priori state estimate with measured data by

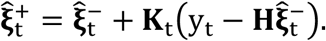

The weighting to correct the a priori augmented state estimate, **K**_t_, is known as the Kalman gain^41, 62^. The Kalman gain is calculated using the available information regarding the confidence in a prediction of the augmented states through the model and the observation model that includes noise by

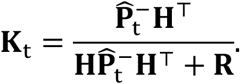

The a posteriori state estimate covariance is then updated by using the Kalman gain to provide the correction

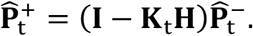

After each time step, the a posteriori estimate becomes the prior distribution for the next time step, and the filter continues.

The Kalman filter requires 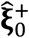 and 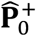 to be initialised to provide the a posteriori state estimate and state estimation covariance for time *t* = 0. We initialised 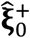 and 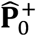 by calculating the mean and covariance of the augmented state vector with the second half of the forward simulation using the default parameters known to generate alpha rhythms^20, 35^, to obtain a stable model output.

The Appendix further explains how the mean and covariance of the augmented state can be estimated for fully nonlinear NMMs.

### Strong and weak posterior alpha power-based contrast imaging

It is known that when people are resting, the alpha oscillations at the back of the brain are very strong, especially in the visual cortex and the surrounding occipital region^10-12, 30^. To study the link between neurophysiological mechanisms and resting-state alpha oscillations, the whole brain MEG-derived source recordings were band-pass filtered in the frequency range of 8-12 Hz to extract the alpha power time-series. The alpha band of this group of participants was determined by the dominant alpha frequency of each MEG source point for all participants, and it was found that more than 90% of the MEG source points had an alpha peak frequency in the range of 8-12 Hz. The alpha power envelope measures the amplitude of alpha oscillations and was used as a research target. We calculated the root-mean-square envelope with a sliding window of length 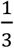 second to indicate the temporal evolution of alpha power. In this study, a labelling scheme was defined to identify strong and weak alpha oscillations in the occipital part of the brain. For each subject, the strong and weak occipital alpha power thresholds were determined by the aggregate averaged alpha power of the source points in the occipital lobe. When the posterior alpha power in the occipital lobe exceeds the 75^th^ percentile of the spatially averaged time-series, the relevant time periods were labeled as strong alpha. When the posterior alpha power is less than the 25^th^ percentile of the spatially averaged time-series, the relevant time periods were labelled as weak alpha. Then, the time indices of strong and weak posterior alpha were respectively used to extract the neurophysiological variable estimates. In this way, two sets of neurophysiological variable estimates were obtained for the whole brain MEG source points related to strong and weak posterior alpha oscillation.

For each subject, a two-sample t-test was performed to compare the sample mean of the neurophysiological variable estimates between the strong and weak occipital alpha in each MEG source point. Such that, for every neurophysiological variable, there were 4714 t-statistics to reflect the sample mean differences across the whole brain. Here let ***X*** and ***Y*** be two random variables, *m* and *n* are the sample sizes for ***X*** and ***Y*** respectively, the two-sample t-statistic is defined as

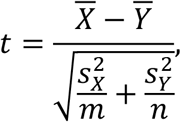

where 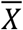 and 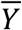 are the sample means, 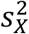 and 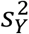 are the sample standard deviations. In our study, ***X*** and ***Y*** respectively represented the neurophysiological variable estimates with respect to strong and weak occipital alpha for each source point. The number of degrees of freedom is given by Satterthwaite’s approximation and calculated by^65, 66^

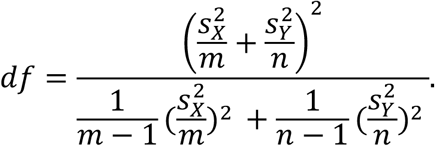

### Pearson’s correlation imaging

Pearson correlation coefficient, ***ρ***_*X,Y*,_ is a measure of linear correlation between two random variables. It is the ratio between the covariance of two random variables and the product of their standard deviations, i.e., for a population, here let ***X*** and ***Y*** be two random variables:

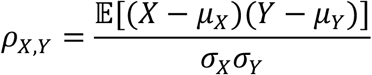

where *µ*_*X*_ and *µ*_*Y*_ are the mean of ***X*** and ***Y***, respectively, and ***σ***_*X*_ and ***σ***_*Y*_ are the standard deviation of ***X*** and ***Y***, respectively.

To measure the linear correlation between the neurophysiological estimate time-series and the alpha power time-series, for each subject sample Pearson’s correlation coefficient, ***r***_*XY*_, was calculated in each source point. ***r***_*XY*_ is calculated by

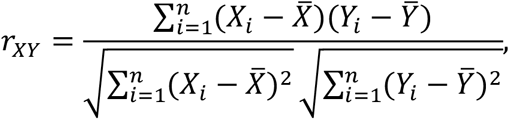

where ***Y***_*i*_ is alpha power envelope time-series, 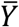 is the mean of the alpha power envelope, ***X***_*i*_ is a given neurophysiological variable estimates’ time-series, and 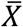 is the mean of the neurophysiological variable estimates.

### Statistical analysis for group-level whole-brain imaging

The group-level analysis was designed to provide a general pattern of the neurophysiological mechanisms for the resting-state alpha rhythm, which merged statistics from individuals. The whole-brain imaging presented a statistical image: at each voxel, a statistic of interest was computed. The group-level whole-brain imaging showed group-level significant voxels for the statistic of interest, and in this study, two statistics were considered as mentioned above: two-sample t-statistic (strong and weak alpha contrast imaging) and correlation coefficients. For one type of statistic, every source point had 22 statistic values derived from 22 subjects. It is reasonable to assume these statistic values were independent and identically distributed random variables regardless of how they were calculated, since for every source point the statistic value was calculated independently from 22 randomly selected participants. To merge statistics from 22 subjects for every source point, a group-level t-statistic was derived by normalising the mean of them by the sample standard deviation. As the whole-brain statistic imaging is a voxel-by-voxel hypothesis testing framework, the multiple comparisons problem ought to be handled. This study employed a nonparametric permutation approach to correct the group-level t-statistic^25^, which provided results similar to those obtained from a comparable statistical parametric mapping approach using a general linear model with multiple comparisons corrections derived from random field theory^67-69^.

With the nonparametric permutation test, the group-level t-statistic image was thresholded at a given critical threshold, and the voxels with group-level t-statistic values exceeding the threshold had their null hypothesis rejected. Rejection of the omnibus hypothesis (that all the voxel null hypotheses are true) happens if any voxel group-level t-statistic value surpasses the threshold, a situation clearly determined by the maximum value of the group-level t-statistic image over the whole brain. Therefore, consideration of the maximum group-level t-statistic handles the multiple comparisons problem. For a valid omnibus hypothesis, the critical threshold makes the probability that the maximum group-level t-statistic exceeds it less than the significance level (*α* = 0.05). Therefore, the distribution of the maximal of the null group-level t-statistic image is needed. The nonparametric permutation approach was employed to yield the distribution of the maximal group-level t-statistic over the whole brain. Since the two individual-level statistics used in this study, the two-sample t-statistic (strong and weak alpha contrast imaging) and correlation coefficients, can be positive or negative, the related null hypothesis is that the group-level t-statistic is zero (i.e., indicating there is no significant difference of the neurophysiological variable estimates between strong and weak alpha for the t-statistic, or no linear correlation between neurophysiological variables and the alpha power for the correlation coefficient) and the alternative is inequality, which is suitable for a two-sided permutation test. The permutation was performed based on the null hypothesis and in this study, for each source point individual-level statistic values from all subjects were multiplied by +1 or -1. Then group-level t-statistic was derived with the permuted statistics by finding the mean of the individual-level statistic and normalised by the sample standard deviation. The maximal group-level t-statistic was noted across the whole brain (4714 source points). The permutation was repeated for 5000 times, resulting in a permutation distribution of the maximal group-level t-statistic given the omnibus hypothesis was true. We can reject the null hypothesis at any source point if the original group-level t-statistic (without permutation) is in the top 100*α* % of the permutation distribution for the maximal group-level t-statistic. In the two-sided permutation test, the critical threshold is 97.5 percentiles and 2.5 percentiles of the permutation distribution for the maximal group-level t-statistic. Then the null hypothesis at any source point can be rejected at the group level with the original t-statistic value (without permutation) exceeding the critical threshold.

With MATLAB package Fieldtrip and customized scripts, the whole-brain imaging was generated by projecting significant group-level t-statistics (after corrections for multiple comparisons) to the corresponding cerebrocortical source points on the template MRI. The imaging was further rendered by linearly interpolating the value of each source point to the surrounding cerebrocortical voxels^51^. Then AAL atlas labels^56^ of the brain ROIs where significant source points were located were noted. Consequently, the group-level whole-brain imaging showed the statistically significant brain areas with respect to the statistic of interest (e.g., strong and weak occipital alpha two-sample t-test, Pearson’s correlation between local alpha power and each neurophysiological variable estimate).

## Results

### Individual example of whole-brain imaging of resting-state neural mechanisms

To illustrate the approach for one individual, **Fig. 1b** shows a sample MEG source time-series from an occipital cortical source (MNI-coordinates: 12, -90, 24) derived by the linearly constrained minimum variance (LCMV) beamformer^16^. The figure focuses on occipital cortex because it is known to be a strong generator of resting-state alpha rhythm. **Fig. 1c** shows the extracted alpha oscillation in red and its power envelope in green. **Fig. 1d-k** illustrates the corresponding estimates of the neurophysiological variables of the NMM, obtained by the AKF. The estimates of the variables converge after a short time and are relatively stable during the resting-state. Moreover, the time-resolved variable estimates afford the possibility to analyse their relationship with other target variables whether they are endogenous or exogeneous in nature. Here for simplicity their relationship with the intermittent waxing and waning of alpha rhythm power (root-mean-square envelope) during resting was considered. This was analysed further at the group level in the following sections, however, as a first step **Fig. 1I** displays example whole-brain contrast imaging results for the same individual in **Fig. 1b-k** where the neurophysiological variable estimates of each source point obtained during strong and weak occipital alpha are contrasted using a two-sample t-test (*α* = 0.05). Only significant voxels were activated after multiple comparisons correction for the whole brain. Strong and weak alpha were defined when the occipital alpha power was greater than the 75^th^ percentile and less than the 25^th^ percentile, respectively. Then the strong and weak alpha time indices were used to extract the corresponding neurophysiological variable estimates. Note only the neurophysiological variable sub-images should be interpreted with respect to statistical testing, not the alpha power sub-image on the left because the null hypothesis is that a given variable is equal during both high and low alpha power periods. Application of the t-statistic to the alpha power sub-image was done purely as a standardised way to view the data. It can be seen that increases in the cortical input, *µ*, is a strong determinant of increased posterior alpha power, while the other neurophysiological variables also appear to contribute. It is also important to note that the NMM only approximates population activity in the cerebral cortex. Therefore, associated variable estimates should be considered approximations of true neurophysiological changes.

A comparison (**Supplementary Fig. 1-4**) shows the superiority, in this context, of the AKF over the most common Kalman filtering scheme for nonlinear systems, unscented Kalman filtering (UKF). Briefly, the results indicated that the AKF is 7.5 times faster than the UKF in processing 3.75-minutes of data, 10% more accurate in tracking model variables, and has similar convergence time. For tracking the actual measurement, AKF showed excellent accuracy in both time and frequency domain (**Supplementary Fig. 5** and **6**).

### Group level whole-brain imaging of resting-state neural mechanisms

The same individual analysis presented in **Figure 1** was applied to MEG resting-state data from 22 subjects and a group analysis was subsequently performed. There was 3.75 minutes of data per subject after pre-processing and artefact removal. Along with traditional contrast-based imaging^24^, this study provides correlation-based imaging as an alternative type of visualization that leverages the space-time resolved nature of the neurophysiological variable estimates to illustrate the neural mechanisms of modulating alpha power. Contrast imaging revealed the mean difference of the neurophysiological variables inferred during strong and weak occipital alpha power. Whereas the linear correlation between neurophysiological variables and associated local alpha power was uncovered by Pearson’s correlation. For each type of imaging and each source point, statistics were derived first at the individual level, and finally merged into the group-level with a t-statistic by normalising the mean of statistics from all subjects by the sample standard deviation. The multiple comparisons problem was handled by a nonparametric permutation approach^25^. The six most significant brain structures associated with each neurophysiological variable were noted in **Supplementary Table 1** and are discussed further below. The max t-statistic and the voxel’s coordinate were also presented.

**Contrast imaging Figure 2** shows the group-level contrast imaging of the mean difference of alpha power and neurophysiological variable estimates between strong and weak occipital alpha power. As with **Fig. 1l**, only the neurophysiological variable sub-images should be interpreted with respect to statistical testing.

**Figure 2.**
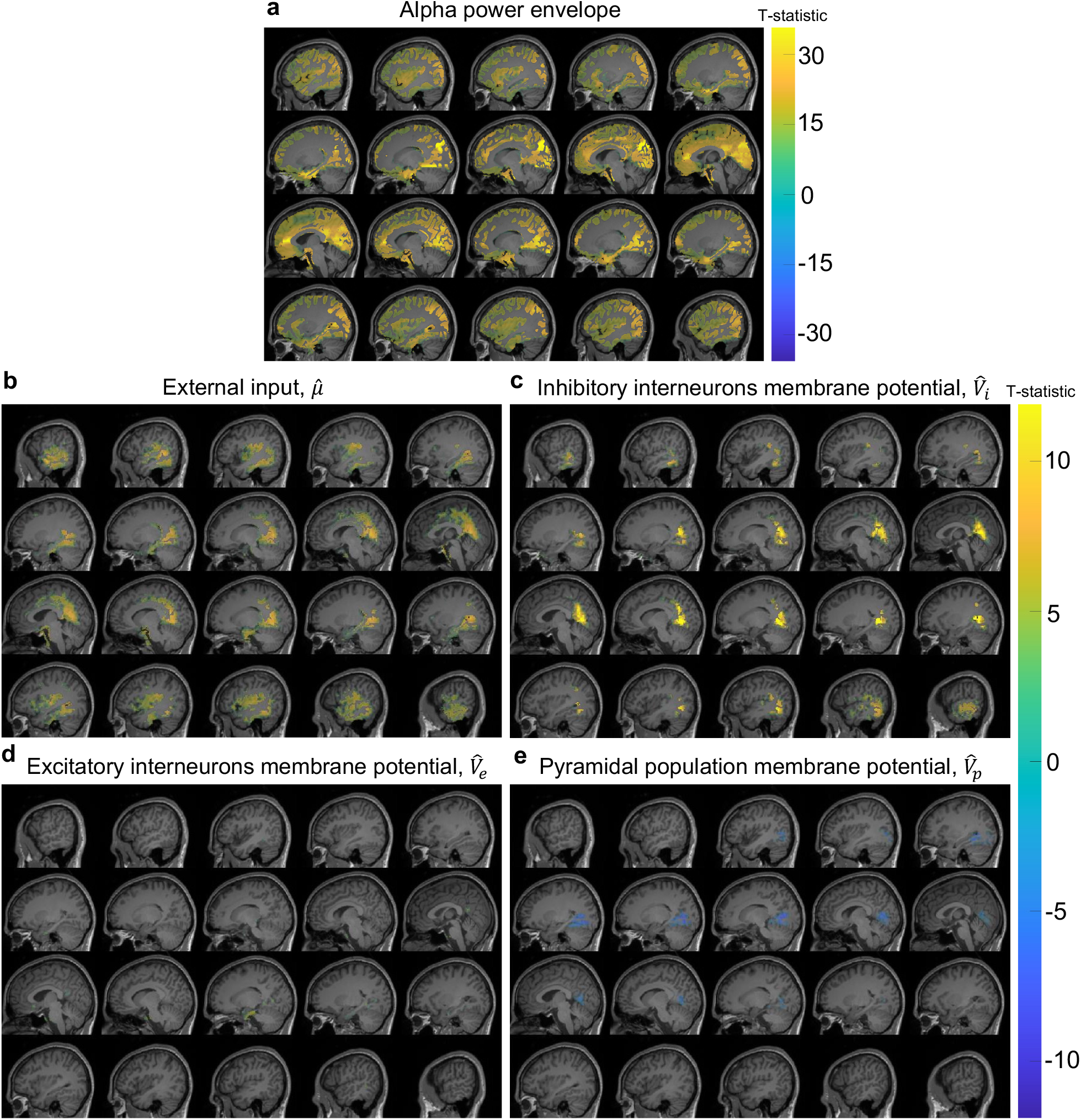
Group-level whole-brain contrast imaging shows the mean difference of alpha power (rms envelope) and neurophysiological variable estimates between two conditions: occipital strong and weak alpha oscillations. For each source point, a two-sample t-test was performed to evaluate the significance of the mean difference of the neurophysiological variable estimates between the aforementioned two conditions. The activated areas in the sub-images represents statistically significant difference was observed after corrections for multiple comparisons (significance level *α* = 0.05). See methods for more details about the statistical analysis scheme. (**a**) Contrast imaging of alpha power. (**b**) – (**e**) Contrast imaging of external cortical input, the mean inhibitory interneuron membrane potential, the mean excitatory interneuron membrane potential, and the mean pyramidal membrane potential. Only neurophysiological variables that showed statistically significant changes after correction for multiple comparisons are displayed.

In **Fig. 2a**, the brain exhibited strong bilateral posterior alpha oscillations. The alpha oscillation in the occipital lobe was the most noticeable, whereas the parietal and temporal lobes had weaker alpha oscillations. Bilateral cingulate gyrus also showed a clear alpha rhythm. The medial cerebral cortex showed more potent alpha oscillations than the lateral. Across the whole brain, four hotspots can be identified from **Fig. 2a**: right cuneus (MNI-coordinates: 12, -90, 24, group-level t-statistic≈30), right calcarine fissure (MNI-coordinates: 12, -96, -6, group-level t-statistic ≈ 29), right anterior cingulate gyrus (MNI-coordinates: 6, 36, 12, group-level t-statistic ≈ 27) and right precuneus (MNI-coordinates: 6, -48, 18, group-level t-statistic≈26). The localised relationship between alpha power and each neurophysiological variable estimate for the identified brain regions is illustrated in **Supplementary Fig. 7. Fig. 2b** shows that strong occipital alpha power was primarily related to large external input, *µ*, in the bilateral occipital and temporal lobes and particularly in the right middle temporal gyrus, bilateral calcarine fissure and surrounding cortex, and bilateral lingual gyrus. In **Fig. 2c**, occipital alpha power change was strongly related to the mean inhibitory interneuron membrane potential, *V*_*i*_, in the left lingual gyrus, bilateral precuneus, left cuneus, and bilateral calcarine fissure and surrounding cortex, where high *V*_*i*_ caused strong occipital alpha power. **Fig. 2d** shows only a few illuminated voxels in the occipital and temporal lobes, implying that the occipital alpha power change has a trivial relationship with the excitatory interneurons. In **Fig. 2e**, the pyramidal membrane potential, *V*_*p*_, at the back of the head dropped when the occipital alpha power was strong and especially in the right lingual gyrus, fusiform gyrus, inferior occipital gyrus, calcarine fissure and surrounding cortex, and inferior and middle temporal gyri.

### Pearson’s correlation imaging

Pearson’s correlation imaging in **Figure 3** shows the brain areas with significant group-level linear correlation between each neurophysiological variable and local, rather than posterior, alpha power.

**Figure 3.**
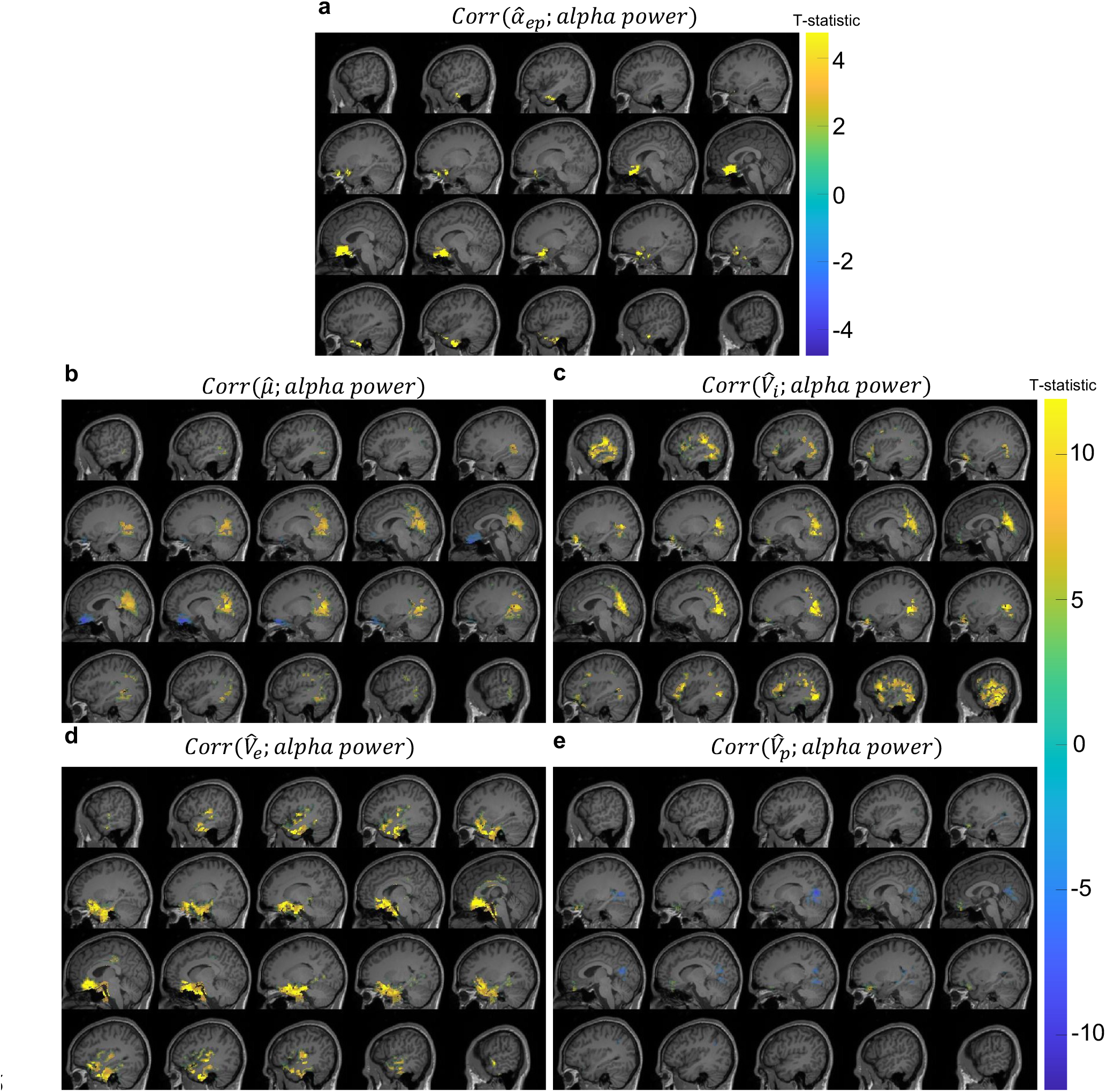
Group-level whole-brain Pearson’s correlation coefficient imaging reveals brain regions where a significant linear correlation between neurophysiological variable estimates and alpha power was observed. For each source point, a one-sample t-test was performed to determine whether the mean of Pearson’s correlation coefficients across all subjects is different from zero. The activated areas in the sub-images represents statistically significant difference was observed after corrections for multiple comparisons (significance level *α* = 0.05). See methods for more details about the statistical analysis scheme. (**a**) – (**e**) Correlation imaging between alpha power and the excitatory to pyramidal connection strength, external cortical input, the mean inhibitory interneuron membrane potential, the mean excitatory interneuron membrane potential, and the mean pyramidal membrane potential. Only neurophysiological variables that showed statistically significant changes after correction for multiple comparisons are displayed.

In **Fig. 3a**, the mean excitatory to pyramidal connection strength, *α*_*ep*_, was linearly related to alpha power in bilateral frontal lobes, and especially in the right medial orbital part of superior frontal gyrus, bilateral gyrus rectus, left olfactory cortex, and left orbital part of superior frontal gyrus. In **Fig. 3b**, the external input, *µ*, was correlated with alpha power in bilateral occipital and frontal lobes. Particularly, positive correlation was observed in bilateral calcarine fissure and surrounding cortex, and left lingual gyrus, while negative correlation was observed in the left gyrus rectus and left orbital part of superior frontal gyrus. **Fig. 3c** shows that the alpha power exhibited positive correlation with the mean membrane potential of inhibitory interneurons, *V*_*i*_, in bilateral occipital and temporal lobes. Left lingual gyrus, left inferior occipital gyrus, left calcarine fissure and surrounding cortex, left cuneus, and bilateral inferior temporal gyrus were identified as the most prominent brain structures. **Fig. 3d** indicates that the mean membrane potential of excitatory interneurons, *V*_*e*_, had substantial positive correlation with alpha power in bilateral frontal lobes such as left orbital part of inferior frontal gyrus, right medial orbital part of superior frontal gyrus, bilateral gyrus rectus. *V*_*e*_ in the right inferior temporal gyrus and insula cortex also showed significant positive correlation with alpha power. **Fig. 3e** reveals that alpha power increased with the decrease of the mean membrane potential of pyramidal cells, *V*_*p*_, in bilateral lingual gyrus, and bilateral calcarine fissure and surrounding cortex. On the contrary, *V*_*p*_ was positively correlated with alpha power in the frontal lobe such as right medial orbital part of superior frontal gyrus, and right orbital part of inferior frontal gyrus.

### Relationship between resting-state networks and alpha-power-linked neurophysiology

**Table 1** summarises which of the previously well studied resting-state sub-network^26-29^ nodes contained statistically significant alpha-power-linked neurophysiological variable changes for both contrast and correlation imaging. Consistent results are observed for both imaging methods with primarily external input, but also the mean membrane potential variables, having significant and varied influence on alpha power within resting-state sub-network nodes.

**Table 1.**
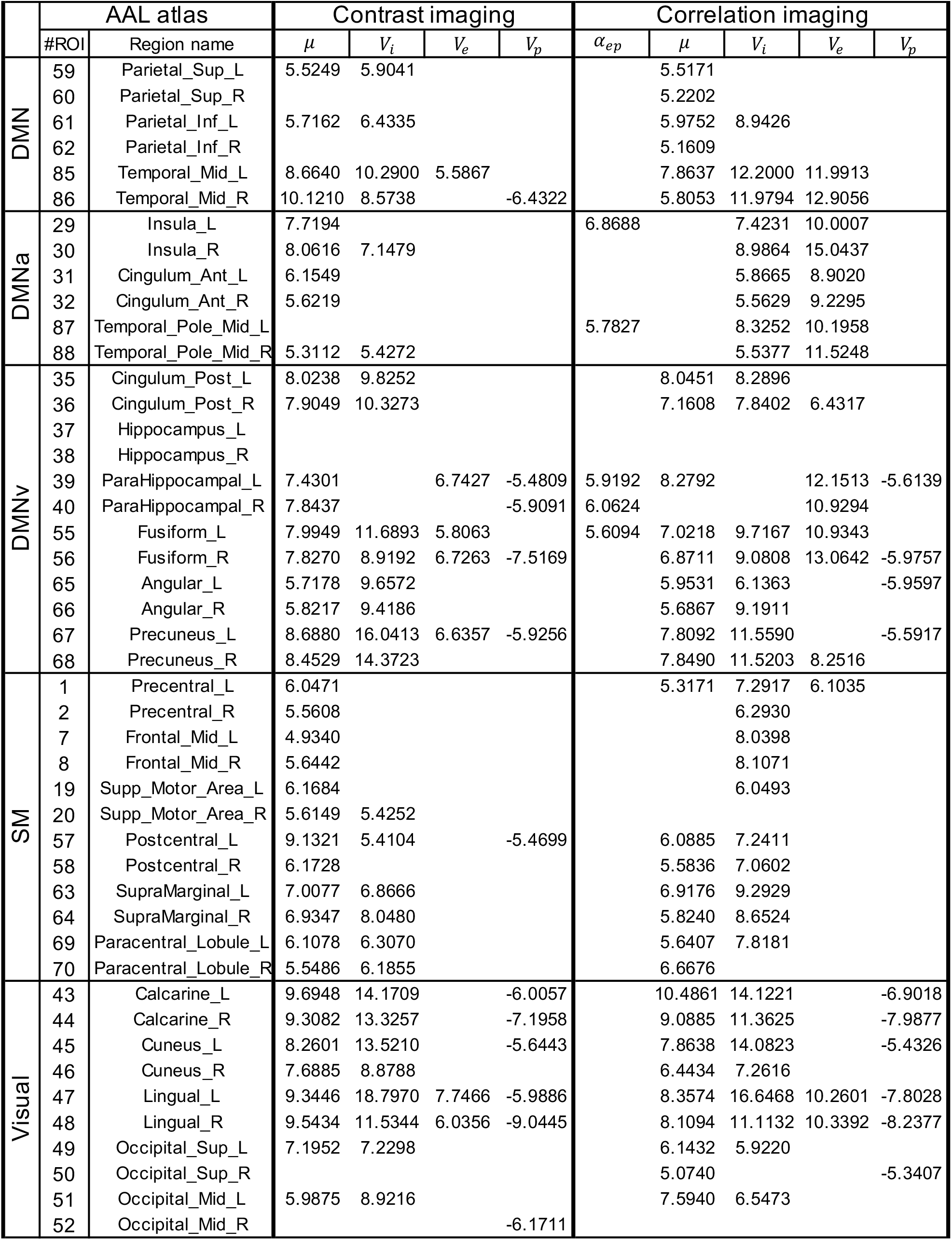
Resting-state sub-networks, their AAL atlas ROIs, and the corresponding maximum group-level t-statistic for statistically significant neurophysiological variables for contrast and correlation imaging. Five resting-state networks are included, namely Default Mode Network (DMN), anterior Default Mode Network (DMNa), ventral Default Mode Network (DMNv), Sensorimotor resting state network (SM), and Visual resting state network (Visual). This table shows if the areas listed in these resting state networks have significant changes in the physiological variables by showing the group-level t-statistic if they are significant, otherwise leaving the space unmarked. Only neurophysiological variables having significant t-statistic source points are presented.

## Discussion

Here a NMM inference-based whole-brain imaging framework has been proposed as a general method to provide reliable spatial and temporal information about the latent neural mechanisms of brain states.

### A novel way to image neural mechanisms across the brain

To quote the famous statistician George Box, this paper takes the perspective that “all models are wrong, but some are useful”. Although the canonical NMM applied here is designed primarily to model alpha rhythms and visual evoked responses^20^, the mathematical framework presented here is general enough to allow researchers to implement arbitrary NMMs that are more relevant and useful for their study of interest. At the same time, many fMRI mapping studies have relied on a canonical HRF^17, 30^, therefore neurophysiological variable mapping studies of the kind presented here that depend on a canonical NMM should also be useful for both imaging and understanding brain function in different ways^9^, despite the obvious approximations of the canonical model. Using more complex and accurate models makes the inference problem more difficult and requires more measurements of brain signals to reduce the non-uniqueness of the solution space and find a more optimal solution. Therefore, the current best path forward is to trade off physiological detail for speed and accurate solutions to achieve results that are useful and interpretable. This is important in human neuroscience as invasive experimental methods typically used to achieve neurophysiological insights are often difficult to apply.

Here the neurophysiological estimates have been visualised with contrast imaging and linear correlation imaging, which associate the neurophysiological variables with a target time-series, alpha-power envelope. However, in general the target could be a behavioural responsiveness variable, exogenous stimulus, or cognitive measure. This suggests broad applicability of the framework. Estimating neurophysiological variables in a time-resolved fashion also opens the door for detecting, and controlling, local critical transitions^31^ across the brain to study different brain states, such as waking, sleeping, intra-awake or other states.

### Neural mechanisms that modulate alpha power

As an example, the framework was applied to resting-state alpha rhythm data to provide new insights. The neural mechanisms underlying the alpha rhythm are still uncertain with debate around the source of the alpha rhythm being the thalamus, thalamocortical loops, the cortex and/or intracortical inhibition^18, 32-34^. Given the complexity of the brain it is likely that alpha rhythms can emerge in different ways depending on axonal and dendritic propagation delays between interacting brain networks or the time constants of excitatory or inhibitory synaptic potentials in these brain networks. Recent human electrophysiology research regarding phase relationships of the alpha rhythm has shown that higher-order posterior cerebral cortex leads lower-order posterior cortex, posterior cortex leads the pulvinar nucleus of the thalamus, and the alpha rhythm is predominantly present in the superficial layers (I/II) of cortex^32^. This suggests a role for the alpha rhythm in top-down feedback processing. While the results presented here for the contrast and correlation imaging, do not focus on which brain areas lead each other, they instead identify the local relationship between alpha rhythm power and population averaged membrane potentials of the pyramidal, excitatory, and inhibitory populations and their population averaged excitatory and inhibitory synaptic connection strengths. This helps to determine which neurophysiological variables are critical contributors to the alpha rhythm across the entire human brain. While phase-lag relationships between the estimated variables could be determined, it is considered beyond the scope of the current paper, and now that the most critical brain areas have been identified in this study, it opens the door for a more focused study that could apply the framework presented here to determine the long-range connection strengths between these critical areas to better determine which areas drive each other. By using the current framework to help narrow down the critical brain areas, it makes the problem of estimating the long-range connection strengths more tractable as it reduces the complexity of the model and the solution space by considering only the critical areas instead of the whole brain.

Regarding the prior observation that alpha rhythm is dominant in the superficial layers of posterior cerebral cortex^32^, the canonical model presented here only includes three neural populations in order to make the inference problem more accurate by keeping it simpler. These pyramidal, excitatory, and inhibitory populations have previously been attributed to the internal pyramidal (V), internal granular (IV), and external granular (II) layers^35^. Consistent with this assignment, the contrast and correlation imaging in **Figure 2** and **3** both revealed that the average membrane potential of the inhibitory interneuron population had a strong positive relationship with alpha power in posterior cortex. This suggests a potential link between the alpha power observed electrophysiologically in superficial layers of posterior cortex and inhibition, however, further work would be needed to study this more accurately using a NMM that considers greater laminar cortical detail.

Regarding the brain areas and networks involved in the resting state and the alpha rhythm, many studies using fMRI and electromagnetic source imaging have identified different resting-state networks^27-29^. With respect to MEG, default mode, somatosensory, visual, and other resting-state networks have been identified with significant hubs tied to frequencies close to alpha in lateral parietal cortex, medial and dorsal prefrontal cortex, and temporal cortex^27, 28^. The contrast and correlation imaging results presented here confirm there are significant changes in the neurophysiological variables, *µ* and *V*_*i*_ predominantly within the previously identified resting-state networks^26-28^, as shown in **Table 1**. Particularly, *µ*, representing cortical input, was critical in resting-state networks showing significant group-level t-statistic in both imaging types. Another interesting observation in the correlation imaging relates to the cortical input, *µ*, which can be regarded as a linear combination of all inputs from all other areas and within the context of dynamical systems theory is known to be a critical bifurcation parameter in the canonical NMM. Within the range of the estimated values of the cortical input, prior mathematical bifurcation analysis of this model showed a u-shaped pattern for alpha power in the model output with escalating *µ*: as *µ* increases, moderate amplitude alpha-like activity emerges in the model, followed by approximately 3 Hz spike-like activity where alpha band content weakens, and finally the model transitions to strong alpha oscillations^36^. **Fig. 3b** demonstrates that *µ* was negatively correlated with alpha power in the medial orbital part of superior frontal gyrus, gyrus rectus and olfactory cortex, and positively correlated with alpha power in the anterior part of occipital lobe. The bifurcation analysis can help explain the inconsistent correlations in frontal and occipital lobes: frontal source points (**Supplementary Fig. 8a-e**) and occipital source points (**Supplementary Fig. 8f-j**) correspond to the left and right sides of the u-shaped relationship between *µ* and alpha power, respectively. Considering *µ* as the inter-regional communication, the result here corroborates earlier EEG-fMRI studies that negative correlation was observed between blood-oxygen-level-dependent functional connectivity and alpha power in the extensive areas of the frontal lobe, while positive correlation was observed in the vicinity of thalamus^37, 38^. More research is needed to investigate the physiological meaning of these correlations.

### Rationality and novelty of the framework

#### Selection of model

The Jansen-Rit (JR) neural mass model^20^ is the canonical model used in the framework. The choice of this model is consistent with the idea of Occam’s razor, that is, one should start with the model with the fewest assumptions that is still able to describe the data. Although many neural models have been specifically designed to describe different brain states in more detail, the model we used is physiologically informative, compatible with M/EEG, and efficient enough for whole brain modelling. Another aspect of the framework is decoupling of the NMMs. This means there are no long-range connections between the NMMs and instead a lumped cortical input is locally estimated. This design allows whole brain modelling and each model fits an MEG source point, 4714 in total, which results in fine-grained neuroimaging with a resolution of 6 mm source spacing. The inference problem for interconnected NMMs would involve estimation of a 4714 × 4714 connection strength matrix and would therefore involve a more complex solution space and require much more computation. Additional constraints are needed to simplify the problem such as using diffusion tensor imaging based tractography^39^ to determine which of the connections should exist, or as noted above once critical brain areas and their neurophysiological changes are identified through the proposed framework, the interconnection of these critical nodes can be investigated in a more tractable study.

#### Innovation of the model inversion scheme

The NMM is a nonlinear dynamical system that maps neurophysiological variables to electromagnetic brain activity. Model inversion techniques estimate unobserved neurophysiological variables in the system from the actual measurement. Bayesian estimation schemes are commonly used in estimating these variables by minimizing the mean squared error. In existing studies, Bayesian inference principally relies on simplifying assumptions, sampling methods, or re-framing the problem for nonlinear systems^40-42^. The extended Kalman filter works on the principle that a linearized transformation of means and covariances of variables of interest are approximately equal to the true nonlinear transformation. Nonetheless, it has been proved that the approximation is not satisfactory and leads to non-optimal estimation^41^. The unscented Kalman filter (UKF) is time-consuming because it reconstructs the mean and covariance by propagating sampled sigma points^41^, and it is not applicable for efficiently estimating variables for thousands of models. The variational Bayes used by DCM iteratively converges to the actual variables, but it is a relatively computationally intensive frequency domain method that cannot effectively provide the evolution of the variables with high time resolution^12, 43-45^. Alternatively, the AKF used in our framework involves a fast, semi-analytic solution to propagate means and covariance of the neurophysiological variables of the fully nonlinear NMM. As indicated in the results the AKF is much more efficient and accurate than the UKF. Importantly, for 3.75 minutes (400 Hz sampling rate) of source time-series data containing 4714 sources in one subject, it only took 3.1 hours to estimate neurophysiological variables from the data using a 6-core CPU 2.9 GHz laptop. Therefore, the framework is reasonably computationally efficient within the context of current neuroimaging methods.

### Framework versatility and modularity

This framework can image neural activity for a wide range of brain oscillations, from delta (1-4 Hz) to gamma (>30 Hz), because the NMM can generate stable output in these frequencies^46^. Moreover, the NMM used in this framework has also been employed in studying the alpha rhythm, epileptiform activity, and anaesthesia in previous works^14, 21, 47^. Therefore, neurophysiological estimates can be used to understand the neural mechanisms behind these brain states. Modularity is another feature of the framework, in which each module is independent and replaceable, i.e., the NMM or even the inference method can be replaced. This approach was taken to enable release of the framework as a software called NeuroMechImager (see Code Availability) so that prior M/EEG source imaging studies could be extended by inputting the source time-series in MNI template coordinates and obtaining neurophysiological variable time-series. This modular approach to the framework for nonlinear neural systems is different from prior end-to-end source imaging work involving only the linear Kalman filter and autoregressive models (also only linear and not neural) which sought to derive brain current sources directly from electromagnetic sensor measurements^48, 49^. The modularity, versatility, efficiency, and accuracy of the framework suggests it will have broad applicability to inferring neural mechanisms underlying different brain states.

## Data and code availability

The 22 healthy male resting-state MEG data and their corresponding MRI data are available at http://consciouscloud.erc.monash.edu (To be made available upon publication). The inference-based whole-brain neural mechanism imaging framework is implemented in a MATLAB package called NeuroMechImager and is available at https://github.com/yundumbledore/NeuroMechImager. A demonstration of the package is provided in the Supplementary Information along with instructions on how to apply the framework to provided demonstration data or other pre-existing source imaging data.

## Acknowledgements

We thank Naotsugu Tsuchiya and David Grayden for helpful preliminary discussions about the work, along with MASSIVE (https://www.massive.org.au) for computational resources. This work was supported by a grant from the Australian Research Council (DP200102600) and Monash University.

## Author contribution

All authors wrote the paper. Experiment planning and data collection involved A.P., S.M., W.W., D.L. and L.K. Development of the inference-based imaging framework and its application involved Y.Z, M.B., P.J.K., D.R.F., Y. L., D.L. and L.K.

## Appendix

As noted above, in general the Kalman filter does not have a solution for nonlinear models. Previous works that used Kalman filtering on the nonlinear NMMs have relied on simplifying assumptions (either linearization of the model, or sampling to estimate the posterior distribution)^47, 63, 64^. This work applied an exact, semi-analytic solution for the mean and covariance of a multivariate Gaussian distribution transformed by the nonlinear NMM. For simplicity of expression, the term B_δ_ξ_t*−1*_∘ ***ϕ***(C_δ_ξ_−1_) can be written as ****ϕ****(ξ_−1_). This solution provides the a priori estimate of the mean,

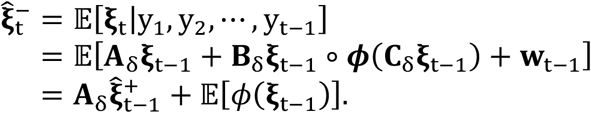

This solution provides the a priori estimate of the covariance,

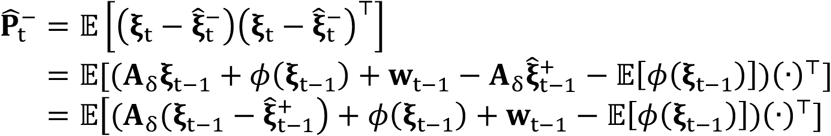

For notational convenience, we denoted the vectors inside the brackets in order a 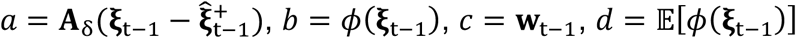, which gives

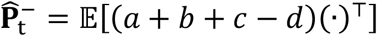

It is given that

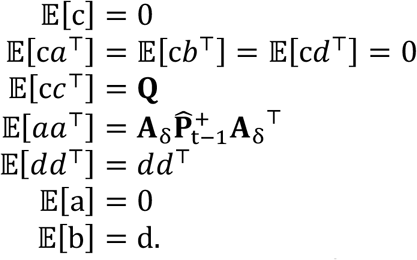

Using the identities to simplify the expression for 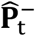 and substituting back in, 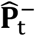 is calculated by

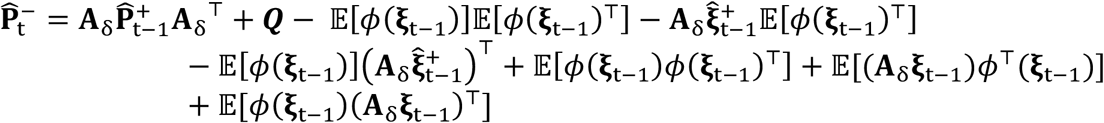

The expectation terms that can be analytically derived are 𝔼{****ϕ****(**ξ**_t −1_)] ,𝔼{(A_δ_**ξ**_t −1_)****ϕ****^T^(**ξ**_t −1_)], and 𝔼{****ϕ****(**ξ**_t −1_)(A_δ_**ξ**_t −1_)^T^] where 𝔼{(A_δ_**ξ**_t −1_)****ϕ****^T^(**ξ**_t −1_)] = 𝔼{****ϕ****(**ξ**_t−1_)(A_δ_**ξ**_t −1_)^T^]^T^. Thus, the analytical solution for 𝔼{****ϕ****(**ξ**_t −1_)] and 𝔼{****ϕ****(**ξ**_t −1_ (A_δ_**ξ**_t −1_)^T^] are derived. A semi-analytical solution is provided for 𝔼{****ϕ****(**ξ**_t −1_)****ϕ****(**ξ**_t −1_)^T^].

For the ease of understanding the following derivations of the above terms, the constant linear terms **A**_δ_, **B**_δ_, **C**_δ_ have been set equivalent to identity matrices of the appropriate dimension. However, in the MATLAB code we used the true matrices and full derivation. Notably, symbols *x*, µ below should not be confused with *x* and ***µ*** above, as we try to follow the convention that *x* is a multi-dimensional random variable and µ is the expectation of the multi-dimensional Gaussian distribution.

For 𝔼{****ϕ****(**ξ**_t −1_)] = 𝔼{**ξ**_t*−1*_∘ ***ϕ***(**ξ**_t −1_)], the expectation of a vector function is a vector and each individual element in the resulting vector can be calculated as below. The simple bivariate case was considered

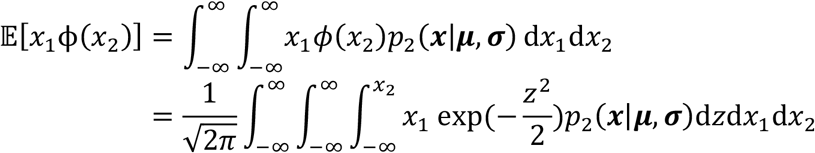

where *p*_2_ is a bivariate Gaussian distribution for *x*_1_, *x*_2_, and 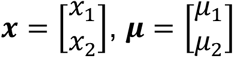 and 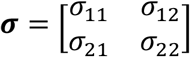 are the expectation and covariance of *x*. We can change the order of integration and shift the *z* integral (using a change of variable to *z*′) giving

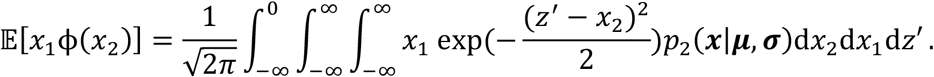

We expand the quadratic inside the exponent of *p*_2_(*x*|µ, *∼*) then collect the *x*_2_ terms inside a single exponent. This gives an integral containing the term of form 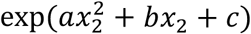, which has the solution

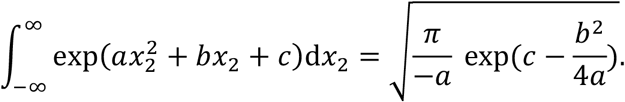

We can solve for the terms *a, b, c* and substitute back into 𝔼{*x*_1_***ϕ***(*x*_2_)] equation, then perform a similar process to collect *x*_)_ terms into the canonical form 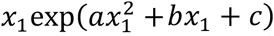. The solution to this second integral is given by

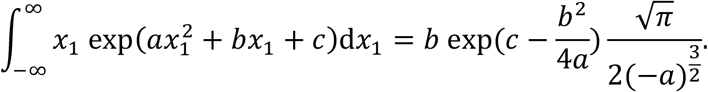

Once we solve for these new *a, b, c* terms and substitute them back into the remaining integral, we can collect the final expression as factors of *z*′ into the form

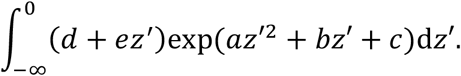

This integral can be solved using a combination of two results. The first is

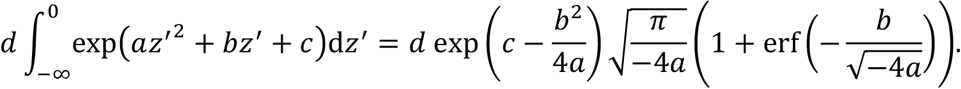

The second solution is

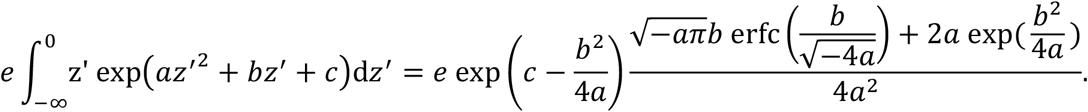

We can combine the above two solutions and simplify to get a general form of the solution for 𝔼{*x*_1_***ϕ***(*x*_2_)],

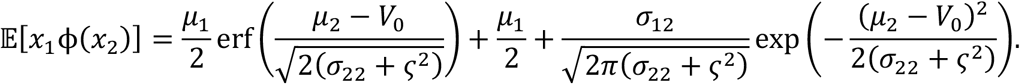

For 𝔼{****ϕ****(**ξ**_t − 1_)(**ξ**_t − 1_)^T^] = 𝔼{**ξ**_t*−1*_∘ ***ϕ***(**ξ**_t − 1_)(**ξ**_t − 1_)^T^], the individual element in the expectation expression can be written into a trivariate form 𝔼{*x*_1_*x*_2_***ϕ***(*x*_*3*_)]. With *x*∼*p*_*3*_(*x*|µ, *∼*),

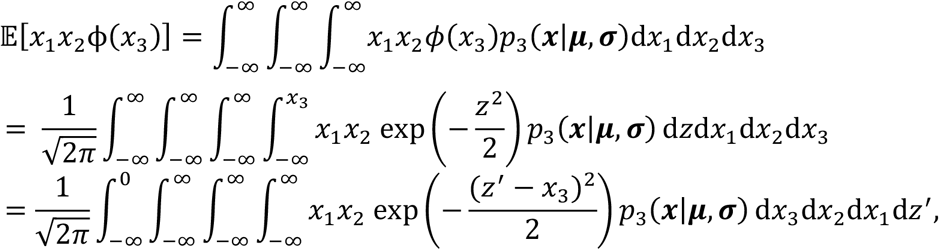

Where 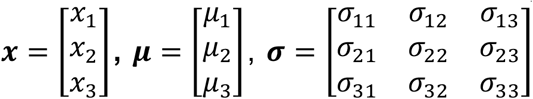. The first integral over *x*_3_ takes the canonical form 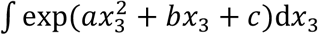, and the second integral over *x*_2_ has a canonical form 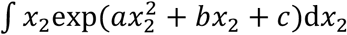. Those two integrals can be solved with previous solutions above. The third integral over *x*_1_ takes the form 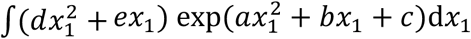, and can be solved by

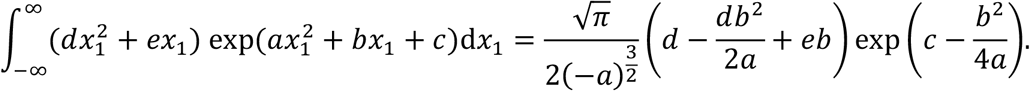

When the coefficients *a-e* are solved and substituted back in the final integral (in *z*′) has the form and solution

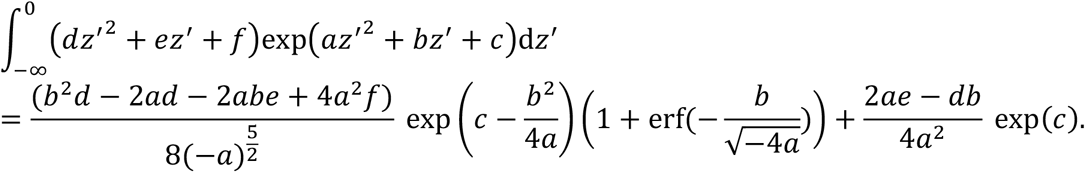

After solving for the coefficients *a-f* and simplifying, the general solution to 𝔼{*x*_1_*x*_2_***ϕ***(*x*_*3*_)] is given by

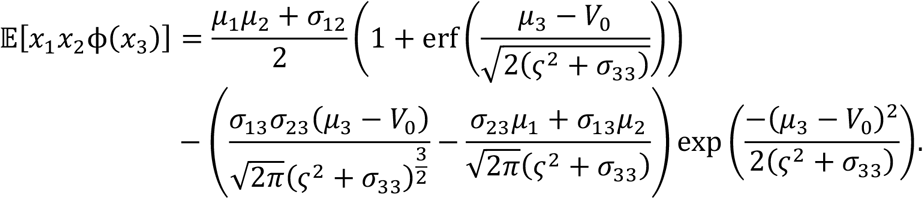

In terms of 𝔼{****ϕ****(**ξ**_t − 1_)****ϕ****(**ξ**_t − 1_)^T^] = 𝔼{(**ξ**_t*−1*_∘ ***ϕ***(**ξ**_t − 1)_)(**ξ**_t*−1*_∘ ***ϕ***(**ξ**_t − 1_)^T^], the terms can be further simplified as 𝔼{***ϕ***(**ξ**_t − 1_)(***ϕ***(**ξ**_t − 1_))^T^ ∘ (**ξ**_t − 1_**ξ**_t − 1_^T^)º. An approximation has been made here 𝔼§***ϕ***(**ξ**_t’)_)(***ϕ***(**ξ**_t’)_)), ∘ (**ξ**_t’)_**ξ**_t’)_,)º = 𝔼{***ϕ***(**ξ**_t − 1_)(***ϕ***(**ξ**_t − 1_))^T^] ∘ (**ξ**_t − 1_**ξ**_t − 1_^T^). The result for the term 𝔼{***ϕ***(**ξ**_t − 1_)(***ϕ***(**ξ**_t − 1_))^T^] can be computed by recognizing that the sigmoid function, ***ϕ***, can be expressed as the Gaussian cumulative distribution function (cdf), with mean *V*_0_ and variance ς. Therefore, the integral can be written as

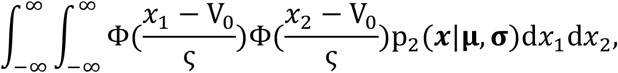

Where 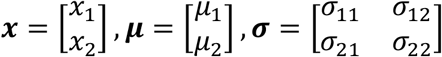. We define two new, independent random variables, *z*_1_ and *z*_2_ that are both normally distributed with mean, V_0_ and variance, ς. As *z*_1_ and *z*_2_ are independent of *x*_1_ and *x*_2_, we have

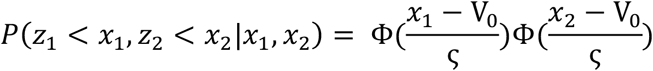

and by the law of total probability, the unconditional distribution is calculated by

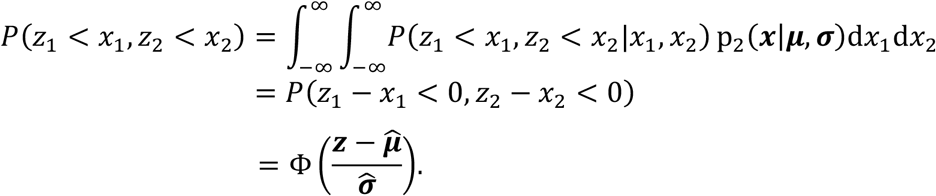

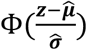 is a bivariate Gaussian cdf, with 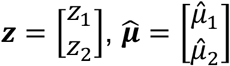, and 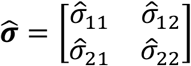. These terms can be calculated by recognizing

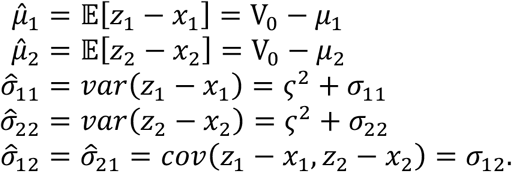

## Supplementary Information

### Supplementary Data

Here we provide additional information to demonstrate the performance of our developed AKF by comparing it to the unscented Kalman Filter (UKF). Then, we provide additional analysis to investigate the relationship between local alpha power and each NMM’s neurophysiological variable estimates for the four identified alpha power hotspots (with the largest group-level t-statistic) in **Fig. 2a** for all 22 subjects. Lastly, we demonstrate NeuroMechImager with two showcases and example data.

### Performance of the Analytic Kalman filter

To demonstrate the performance of the AKF, it was compared with the most commonly used nonlinear Kalman filter, the UKF, with simulated and real experimental data. We conducted controlled variable experiments in which we implemented models in the same way for both filters and initialised them with the same state and covariance. The initial state and covariance were defined by taking the mean and covariance of the augmented state vector from the second half of a 30-second forward simulation with the default model parameters^1, 2^. The main difference between the AKF and UKF is that the AKF is based on a semi-analytic solution to propagate the a posteriori state estimate and covariance, while the UKF applies the unscented transform to enable the propagation. As a result, the UKF is more computationally intensive and less numerically stable/accurate than the AKF.

### Simulated data study

Monte Carlo simulations were performed by testing both filters in the canonical NMM’s parameter space. The artificial data was generated by forward simulating the model with fixed parameters taken from each grid point in the parameter space. We then estimated model states and parameters from the data using both filters. The filter performance was identified for each grid point in the space by calculating data prediction error, parameter estimation error, state estimation error and filter convergence time. We repeated the whole experiment thirty times with a different random seed each time in generating artificial data, and the reported result was the average across all thirty experiments. This was to achieve a reliable and general result, considering that the forward modelling contains Gaussian white noise, and different random seeds result in slightly different artificial data.

The data prediction error was measured by root mean square error (RMSE) between the data prediction and the actual measurement and normalised by the difference between maximum and minimum of the actual measurement, since the actual measurement is a dynamic oscillation. The model state error is an average across four states (i.e., *V*_*ip*_, *V*_*pi*_, *V*_*pe*_, *V*_*ep*_) in the model of the RMSE between states estimates and the ground truth and normalised by the difference between maximum and minimum of the ground truth, since model states are dynamic oscillations. The model parameter error is an average across five parameters (i.e., *µ, α*_*ip*_, *α*_*pi*_, *α*_*pe*_, *α*_*ep*_) in the model of the RMSE between parameters estimates and the ground truth and normalised by the absolute value of the ground truth, since the parameters are held constant during a given simulation. The filter convergence time is defined as the time spent for the last parameter estimate to converge to the ground truth.

The parameter space was defined by positing default values of the neural mass model in the centre of the space and expanding the space until the model cannot generate activity with a dominant spectral peak around the alpha band (8-12 Hz). As we wanted to explore an enormous parameter space, it is safe to put the model default parameters in the middle to maximise the space that can generate expected signals. The space was further confirmed since averaged parameter estimates from the actual data are also within this space. The defined parameter space is shown in **Supplementary Table 2**.

**Supplementary Fig. 1** shows heatmaps of data prediction error (i.e., error between the measured and estimated source signal) of the AKF and UKF. **Supplementary Fig. 1a, b** show that both filters can track the signal with less than 2.5% error. In **Supplementary Fig. 1c**, AKF outperformed UKF in the whole parameter space, and AKF gained 7% less error than UKF. **Supplementary Fig. 2** shows heatmaps of the parameter estimation error of the AKF and UKF. Both filters can accurately converge to the ground truth with less than 6% error. **Supplementary Fig. 2c** shows that AKF is 10% more accurate at tracking parameters than the UKF when averaging across the whole parameter space. Noticeably the UKF exhibited less accuracy when *α*_*ep*_ and *α*_*ep*_ are simultaneously small. **Supplementary Fig. 3** shows heatmaps of model state estimation error, and both filters have a trivial error of less than 2.3%. In **Supplementary Fig. 3c**, AKF is more precise in tracking model states than UKF when averaging across the whole parameter space, and the difference is significant when *α*_*ep*_ is small. **Supplementary Fig. 4** shows heatmaps of filter convergence time. Overall, both filters converge in less than 1 second though UKF has a trivial advantage over AKF. In **Supplementary Fig. 4a**, AKF has a diagonal motif in the heatmap, implying that the three variables shown in the figure are all important in terms of convergence time. For UKF, **Supplementary Fig. 4b** shows the horizontal mode, which means that *α*_*ep*_ is more important in terms of filter convergence time. Specifically, when *α*_*ep*_ is large, the UKF takes more time to converge to the ground truth than AKF.

#### Real data study

Since it is very difficult to measure the ground truth for model states and parameters for real data (hence why this inference method has been developed), an alternative way to validate the filter performance on real data is by evaluating the amplitude and frequency components of the data prediction. The filter works accurately when the data prediction tracks the actual measurement in the time domain and consists of the same components as the actual measurement in the frequency domain. To demonstrate the reliability of AKF on real data, we calculated the data prediction error and coherence between the actual measurement and data prediction while running AKF for each MEG source point and each subject.

The data prediction error was defined as the difference between the data prediction and the actual measurement. **Supplementary Fig. 5** shows the distribution of the data prediction error displayed as probability density functions for each subject to illustrate the normality and whiteness of the error. The probability density function has a narrower peak and heavier tails than a normal distribution with the same mean and variance derived from the actual for each subject. It can be concluded that the AKF was able to track the actual data precisely and the assumptions of Kalman filtering, that the augmented state variables and measurement noise are normally distributed, are approximately valid.

The coherence was calculated to show the correlation in the frequency domain between the data prediction and actual measurement. The coherence is calculated by

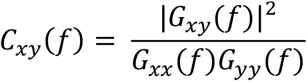

where *G*_*xy*_(*f*) is the cross-spectral density between two signals *x* and *y*, and *G*_*xx*_(*f*) and *G*_*yy*_(*f*) are the power spectral density of *x* and *y*, respectively. The coherence varies between 0 and 1. C_*xy*_(*f*) = 1 means that the actual measurements are perfectly correlated with data prediction at the frequency *f* and C_*xy*_(*f*) = 0 means they are completely uncorrelated. For each of the 4714 MEG source points and each subject, we calculated the coherence between the data prediction and actual measurement and presented the result in **Supplementary Fig. 6**. In each subplot, the red line represents the mean of coherence across 4714 MEG source points and the grey error band refers to its 95% confidence interval. It is noted that the data prediction by AKF is highly correlated with the actual measurement in the frequency band less than 15 Hz. Particularly, in the alpha band 8-12 Hz, the coherence is above 0.75 on average across all subjects. This result suggests that the data prediction by AKF resembles the frequency components in the actual measurement. Together with the result in data prediction error, AKF is a reliable estimator, and its neurophysiological variable estimates can be used to understand the neural mechanisms behind brain states.

Time efficiency is a crucial consideration in the whole-brain imaging framework. The AKF has the inherent advantage of being significantly faster than sampling-based methods such as the UKF. The actual data length in our study is 3.75 minutes per subject, and the data sampling rate is 400 Hz. To quantify the running time for the AKF and UKF, we conducted 100 experiments. Each experiment used the same recording for both filters and different experiments used different recordings. We then noted the time it took to complete the estimation. The results show that the expected running time of AKF is 4.0402 seconds with a standard deviation of 0.0771, and the UKF is 30.7272 seconds with a standard deviation of 0.6923. Taking this statistic into our case study, it only took 529 seconds to finish estimating 4714 time-series with a high performance 36-core CPU workstation for AKF, while UKF is expected to take 4024 seconds.

In summary, AKF is approximately 7.5 times faster than UKF on average and outperforms UKF in estimation accuracy by at least 5%. Besides, AKF enables data prediction highly correlated with the actual measurement in the alpha band, which is of the interest in this paper.

### Localised analysis of the relationship between alpha power and neurophysiological variables

To further study the neurophysiological substrates of alpha power, in **Supplementary Fig. 7** the local relationship between neurophysiological variables and alpha power was visualised for the four identified hotspots in **Fig. 2a** for all 22 subjects. To eliminate individual differences and facilitate direct comparison across individuals, the neurophysiological variable estimates and alpha power have been normalised using student-t statistics.

All four selected areas showed a consistent pattern in terms of the external input, *µ*, that is, when the external input increased, the alpha power increased monotonically. In precuneus and anterior cingulum, alpha power increased because of strong connection strength from inhibitory to pyramidal populations, *α*_*ip*_, considering *α*_*ip*_ is defined negative (strong *α*_*ip*_ means large absolute value of it). Nevertheless, cuneus and calcarine cortex showed conflicting manifestations across individuals for *α*_*ip*_ such that some subjects showed matching patterns whereas other subjects raised the opposite pattern. The mean membrane potential of inhibitory interneurons, *V*_*i*_, exhibited the same physiological modulation for alpha power in precuneus and anterior cingulum, where its increase led to the alpha power first falling and then rising. For the mean membrane potential of the excitatory population, *V*_*e*_, only the precuneus showed a consistent pattern of alpha power across all subjects, where its increase caused alpha power to first fall and then rise.

In summary, the external input, *µ*, and the inhibitory feedback loops including *α*_*ip*_ and *V*_*i*_ have various but critical roles in regulating occipital alpha power. The neural substrates that regulate alpha power vary between subjects and brain regions for the same neurophysiological variable.

### Demonstration with example data

#### Installation guide

The framework is implemented in MATLAB scripts. The framework is most compatible with MATLAB versions equal or later than 2020a. To use the framework, three MATLAB toolboxes are to be installed, namely signal processing toolbox, statistics and machine learning toolbox, and parallel computing toolbox. The imaging module is built upon Fieldtrip (https://www.fieldtriptoolbox.org) and we recommend downloading version 20200828 or later updates. Another link is helpful in adding fieldtrip path to MATLAB (https://www.fieldtriptoolbox.org/faq/should_i_add_fieldtrip_with_all_subdirectories_to_my_matlab_path/). However, note that Fieldtrip is only required for visualisation of the neurophysiological variables, and not for running the neurophysiological variable estimation module. In this demonstration, we used Fieldtrip for visualisation, but one can use other visualisation packages (more information can be found at https://github.com/yundumbledore/NeuroMechImager).

Here the framework is demonstrated via two showcases. The code required to reproduce each case is available at https://github.com/yundumbledore/NeuroMechImager. After one downloads the package, one needs to change the current working directory to this package in MATLAB Command Window and run ‘defaults’, which sets the defaults and configures up the minimal required path settings. The example data consists of 4714 MEG-derived source timeseries in the cerebral cortex from subject 21 and is available to download at https://drive.google.com/drive/folders/18EvFR4cr6YfhNUgZaijj9L3M1sG6cusL?usp=sharing. The data is to be put under the directory ‘/data’. Before running showcases, one needs to make sure that the current directory is ‘NeuroMechImager’. In this directory, one can find a file called ‘test.m’ and this is the script one needs to use to run showcases. Note that there are three critical variables to define before one runs the script:

1. ‘pipeline’: put “AKF estimation” or “Contrast imaging” in the list to indicate which showcase to run.
2. ‘ncpu’: number of CPUs one wants to use. By default, all CPUs will be used.
3. ‘multiple_comparison_correction’: put 0 or 1 to indicate whether to perform corrections for multiple comparisons in Case 2.

#### Case 1: Neurophysiological variable inference by AKF

This showcase demonstrates estimating neurophysiological variables in the neural mass model from 4714 MEG-derived source time-series by the AKF. One needs to open the MATLAB script ‘test.m’ and put “AKF estimation” in the ‘pipeline’ list. One can run the script by entering the following command in the MATLAB Command Window: ‘test’. The input to the framework is subject 21 MEG data containing 4714 virtual timeseries and the output is a (.mat) file containing neurophysiological variable estimates for every source point at ‘/output/variable_estimates_21.mat’. There are eight matrices in ‘/output/variable_estimates_21.mat’ and they are the neurophysiological variable estimates for *α*_*ep*_, *α*_*ip*_, *α*_*pe*_, *α*_*pi*_, *µ, v*_*e*_, *v*_*i*_, *v*_*p*_. Each element contains 4714 rows representing 4714 MEG source points.

### Case 2: Contrast imaging between strong and weak occipital alpha oscillation

This showcase is to image the contrast of neurophysiological variables between when the occipital alpha power is strong and when it is weak. One needs to open the MATLAB script ‘test.m’ and put “Contrast imaging” in the ‘pipeline’ list. The variable ‘multiple_comparison_correction’ toggles between performing corrections for multiple comparisons or not. Note that the correction implemented in this case study is exclusively for individual-level analysis but the principal mechanism is the same with group-level analysis^3^. One can then run the script by entering the following command in the MATLAB Command Window: ‘test’. The input to the framework is the neurophysiological variable estimates derived in Case 1 and the output is imaging showing the contrast of neurophysiological variables during strong and weak occipital alpha power. The output images can be found under the directory ‘/demo_cases/contrast_imaging/visualisation_outputs’. Three dimensions to view the brain are provided: lateral, rear, and dorsal view. **Supplementary Fig. 9** to **Supplementary Fig. 17** show the individual-level (subject 21) contrast imaging in lateral view with corrections for multiple comparisons.

## Supplementary Figures

**Supplementary Figure 1.**
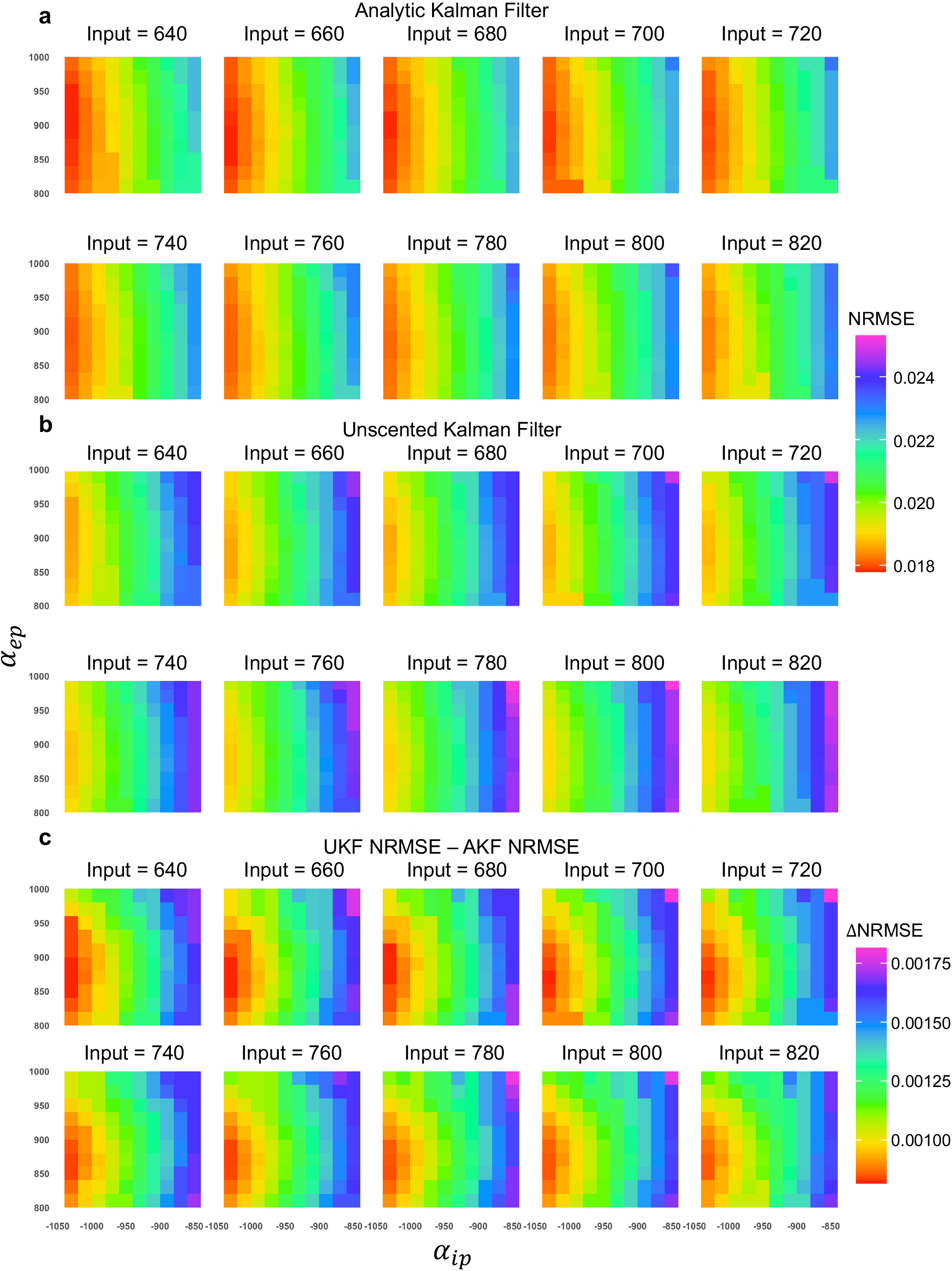
AKF and UKF prediction NRMSE (normalised root mean square error) in a defined parameter space. NRMSE in each tile represents how accurate the filter can track the simulated signal which is generated by forward simulating the model with the fixed parameters’ values taken from the grid point. (**a**) NRMSE of the AKF. (**b**) NRMSE of the UKF. (**c**) NRMSE difference by subtracting AKF NRMSE from UKF NRMSE.

**Supplementary Figure 2.**
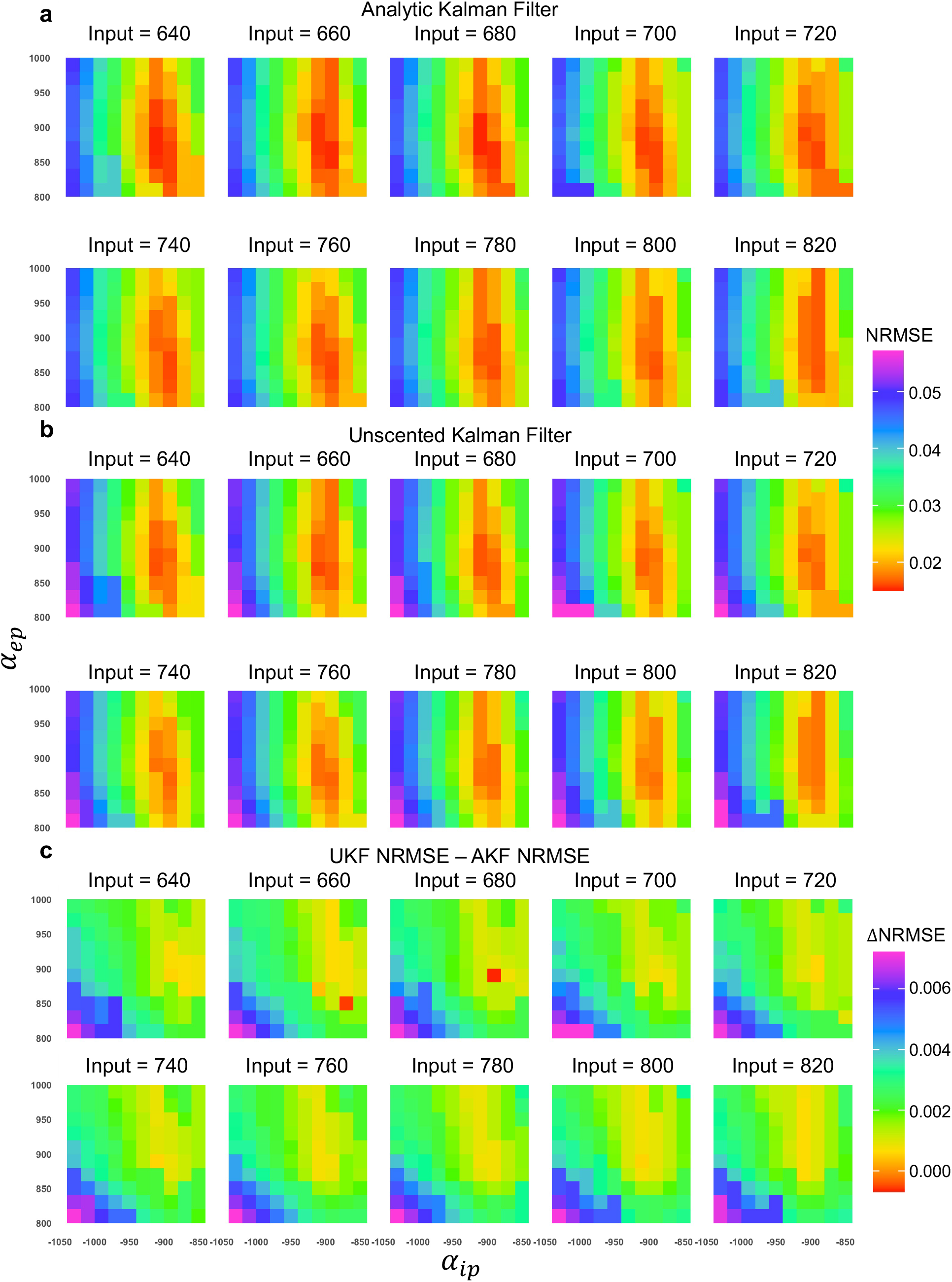
Parameters NRMSE (normalised root mean square error) in a defined parameter space. NRMSE in each tile represents how accurate the filter can track the model parameters given the simulated signal which is generated by forward simulating the model with the fixed parameters’ values taken from the grid point. Since there are five parameters, the NRMSE shown is the average over all of them. (**a**) Parameters NRMSE of the AKF. (**b**) Parameters NRMSE of the UKF. (**c**) Parameters NRMSE difference by subtracting the AKF NRMSE from the UKF NRMSE.

**Supplementary Figure 3.**
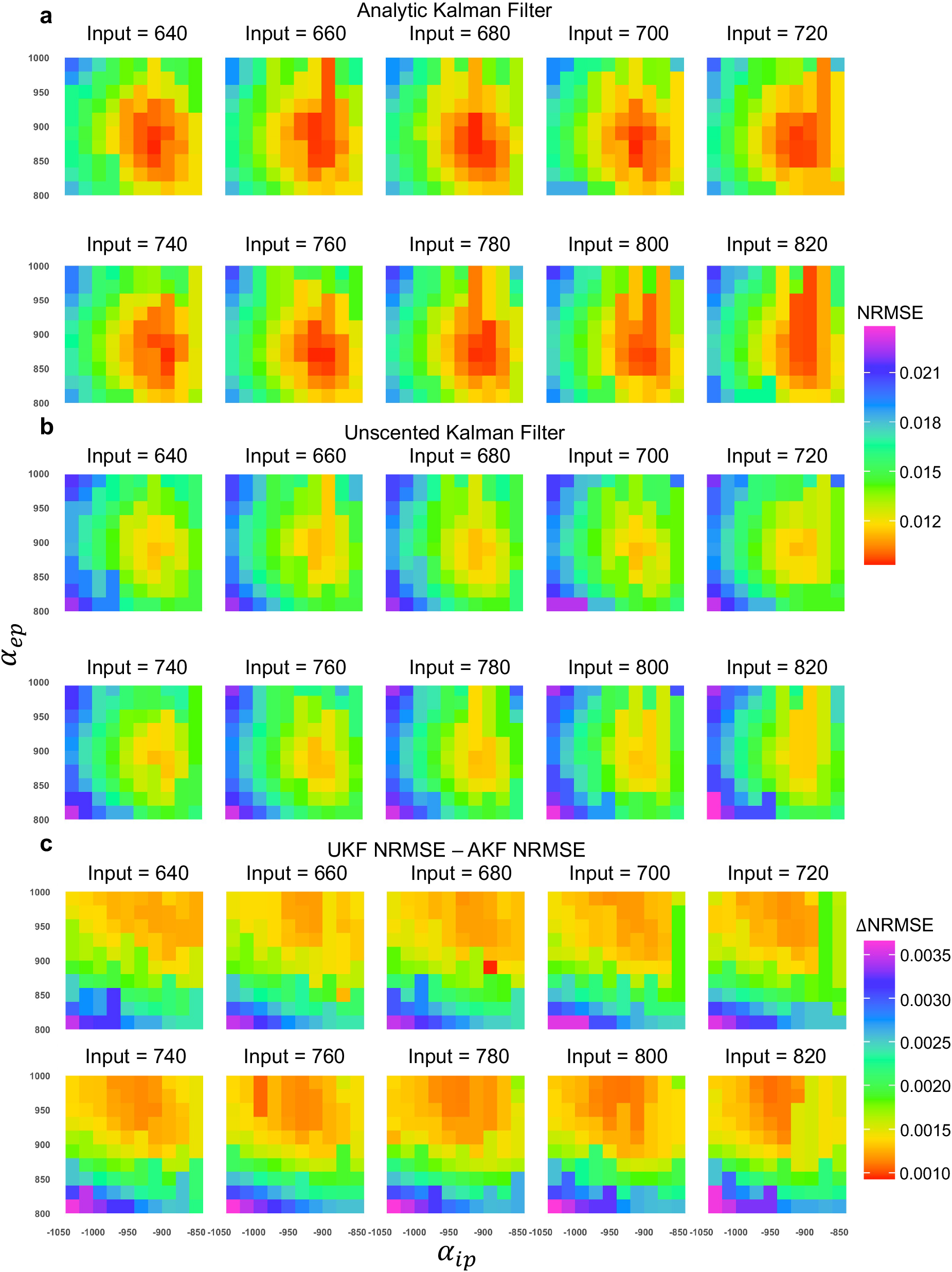
State variables NRMSE (normalised root mean square error) in a defined parameter space. NRMSE in each tile represents how accurate the filter can track model state variables given the simulated signal which is generated by forward simulating the model with the fixed parameters’ values taken from the grid point. Since there are multiple state variables, the NRMSE shown is the averaged over all of them. (**a**) State variables NRMSE of the AKF. (**b**) State variables NRMSE of the UKF. (**c**) State variables NRMSE difference by subtracting AKF NRMSE from UKF NRMSE.

**Supplementary Figure 4.**
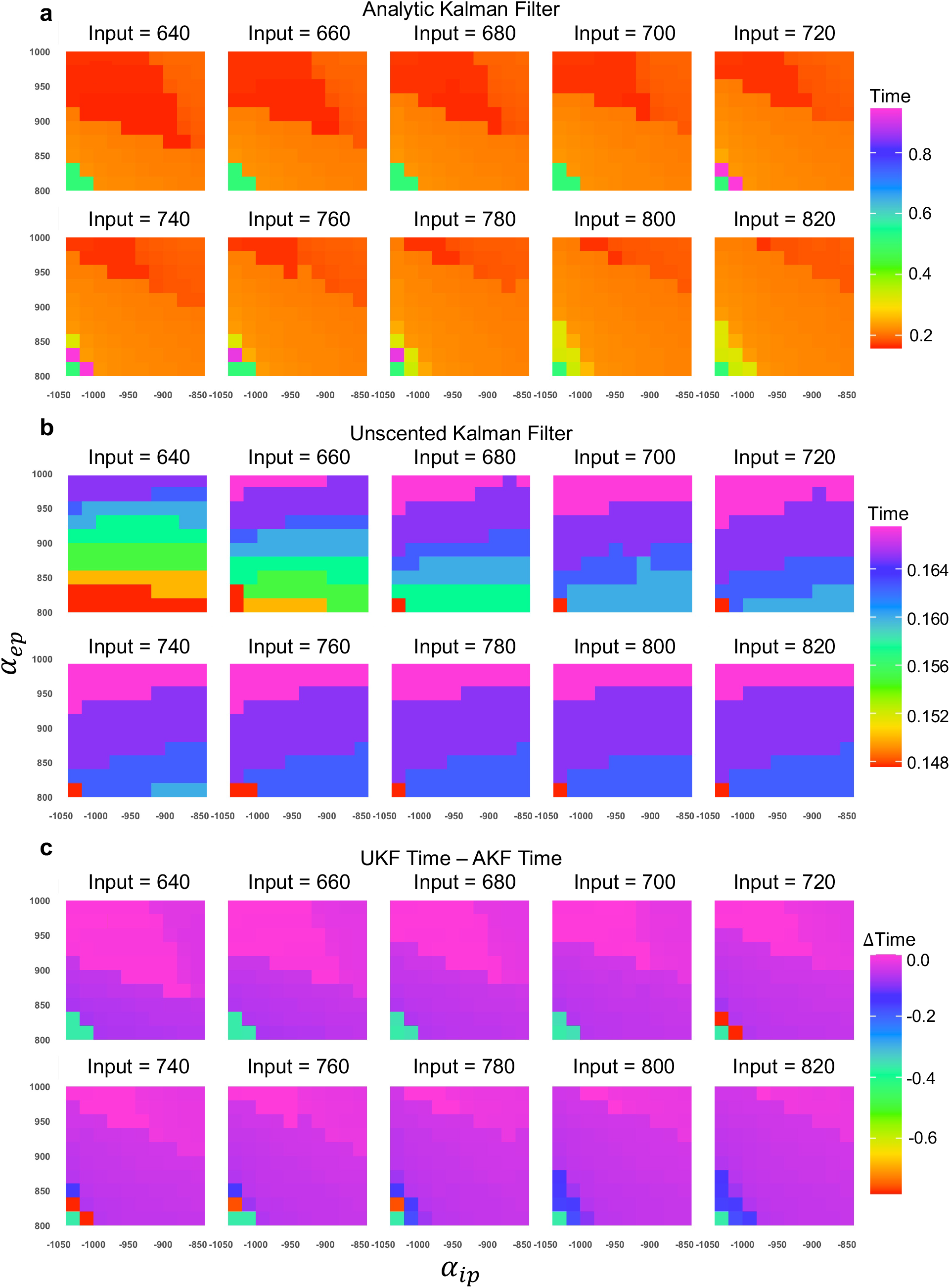
AKF and UKF convergence time in a defined parameter space. The value of each tile represents the time that the last model parameter converges to the ground truth given the simulated signal which is generated by forward simulating the model with the fixed parameters’ values taken from the grid point. (**a**) Convergence time of the AKF. (**b**) Convergence time of the UKF. (**c**) Convergence time difference determined by subtracting AKF convergence time from UKF convergence time.

**Supplementary Figure 5.**
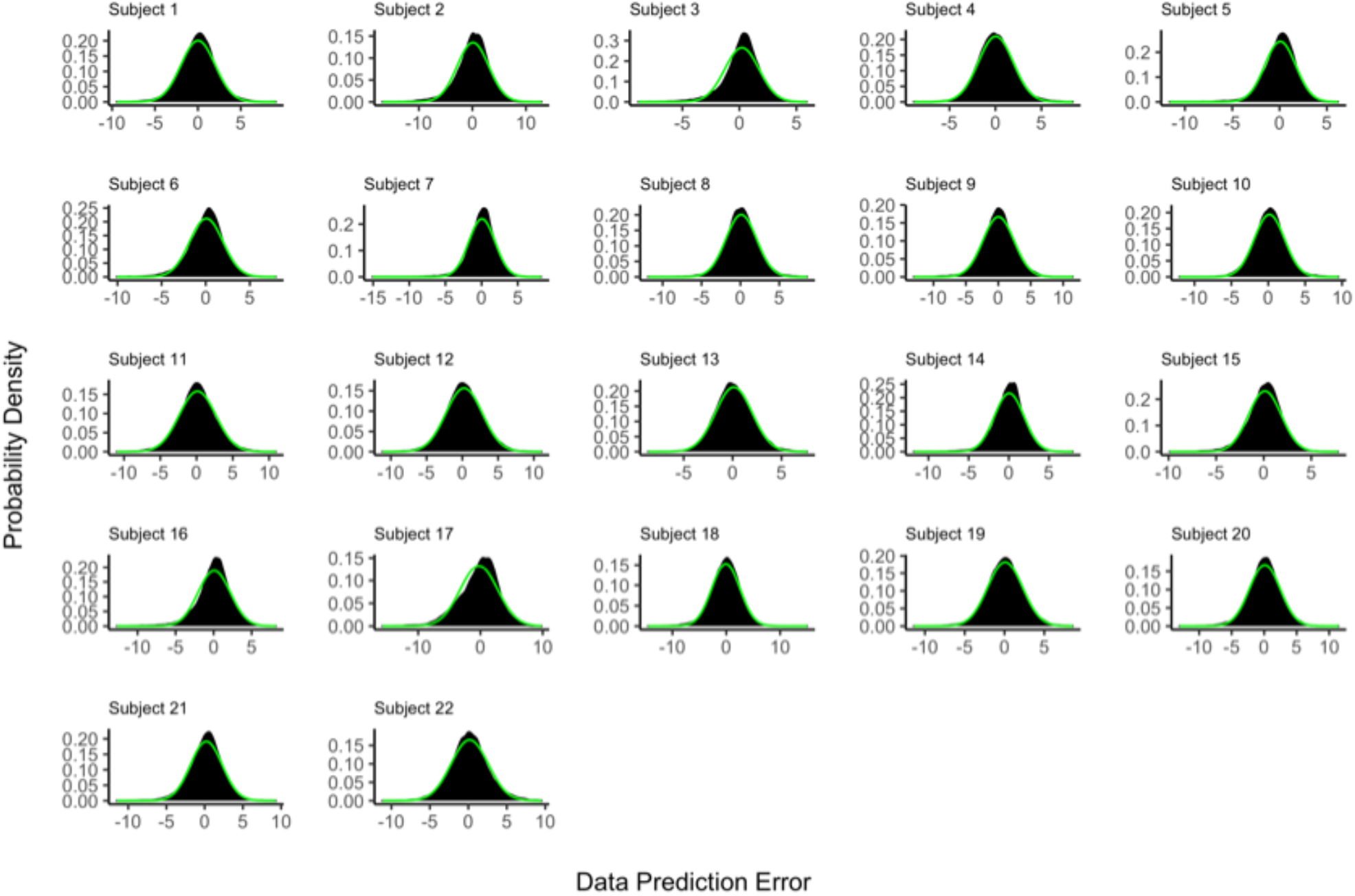
The distributions of the AKF data prediction error (the difference between AKF data prediction and the actual measurement) for all 22 subjects and all 4714 reconstructed MEG source points, displayed as probability density functions (black histogram) plotted against normal probability density functions with the same mean and standard deviation as the data prediction error from each subject (green curves). In each plot, the y-axis represents the value of the probability density, and the x-axis represents the data prediction errors.

**Supplementary Figure 6.**
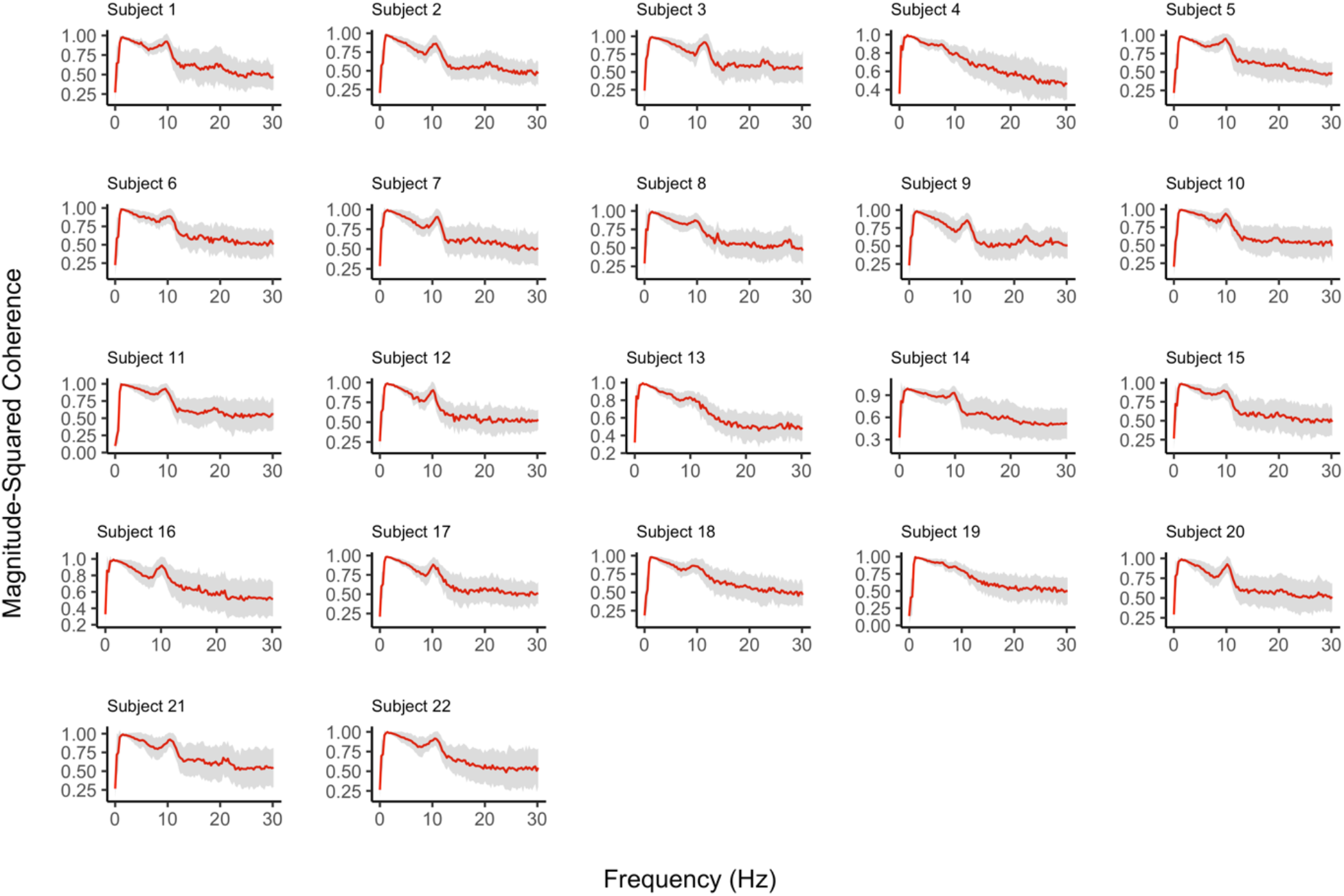
The coherence between AKF data prediction and actual data for all 22 subjects and all 4714 reconstructed MEG source points. Red lines are the mean of the coherence and grey error bands are 95% confidence interval of the mean. Coherence values close to 1 indicates accurate data prediction at the associated frequency.

**Supplementary Figure 7.**
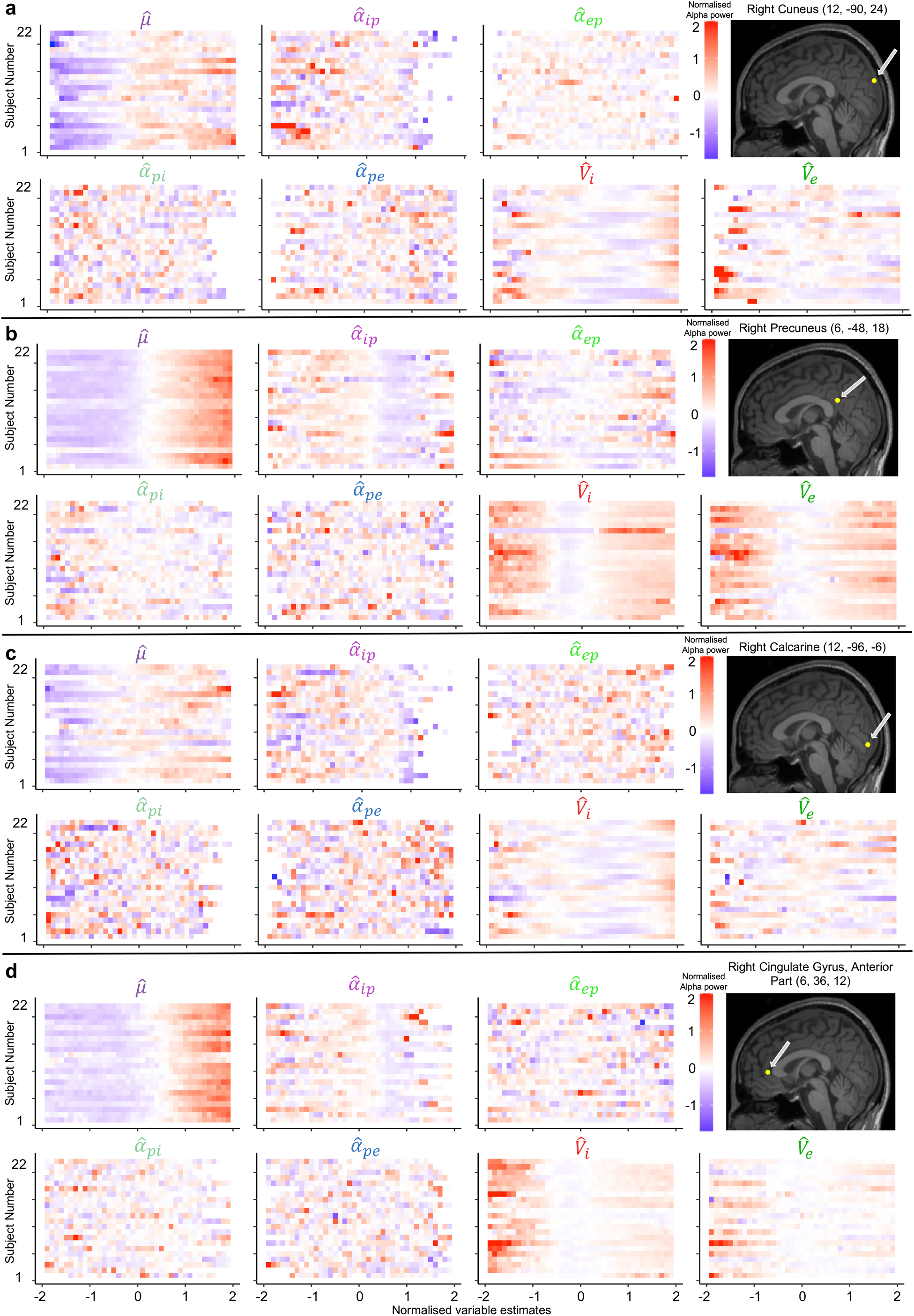
Localised relationship between neurophysiological variable estimates and alpha power for all 22 subjects. Each subpanel is associated with a brain source showing alpha power changes as identified in **Fig. 2a**. Neurophysiological variable estimates and the corresponding alpha power from 22 subjects are normalised and only two standard deviations are shown.

**Supplementary Figure 8.**
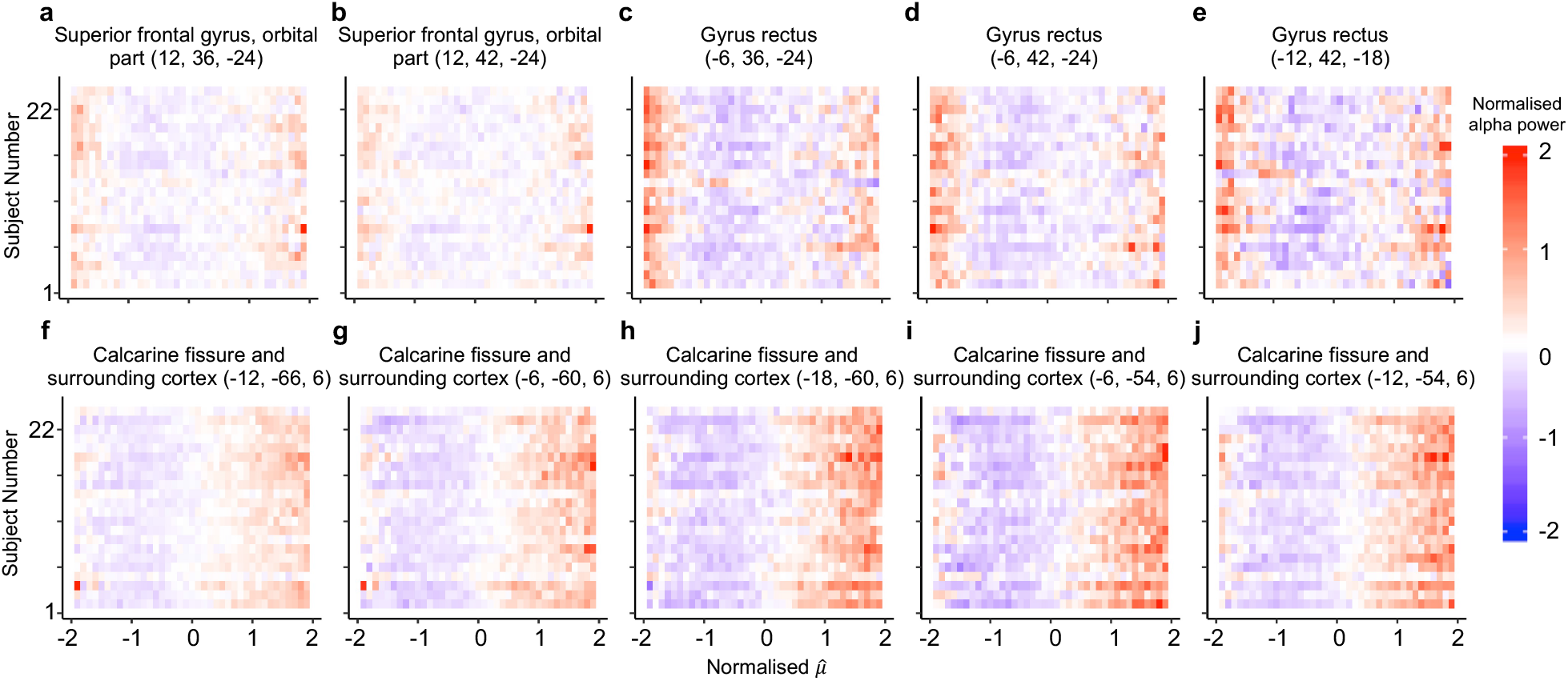
Localised relationship between external input and alpha power for all 22 subjects. Sub-images in the first row are associated with top six source points exhibiting the strongest negative correlation (source points are from the orbital part of superior frontal gyrus and gyrus rectus). A relatively decreasing pattern can be observed, meaning a negative correlation between *µ* and alpha power. The range of 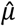 for the identified negative correlation source points can be regarded as the left side of the u-shaped relationship between *µ* and alpha power. Sub-images in the second row are associated with top six source points exhibiting strongest positive correlation (source points are from the occipital lobe). A relatively monotonic increasing pattern can be observed, indicating positive linear correlation. The range of 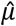 for the identified positive correlation source points can be viewed as the right side of the u-shaped relationship between *µ* and alpha power.

**Supplementary Figure 9.**
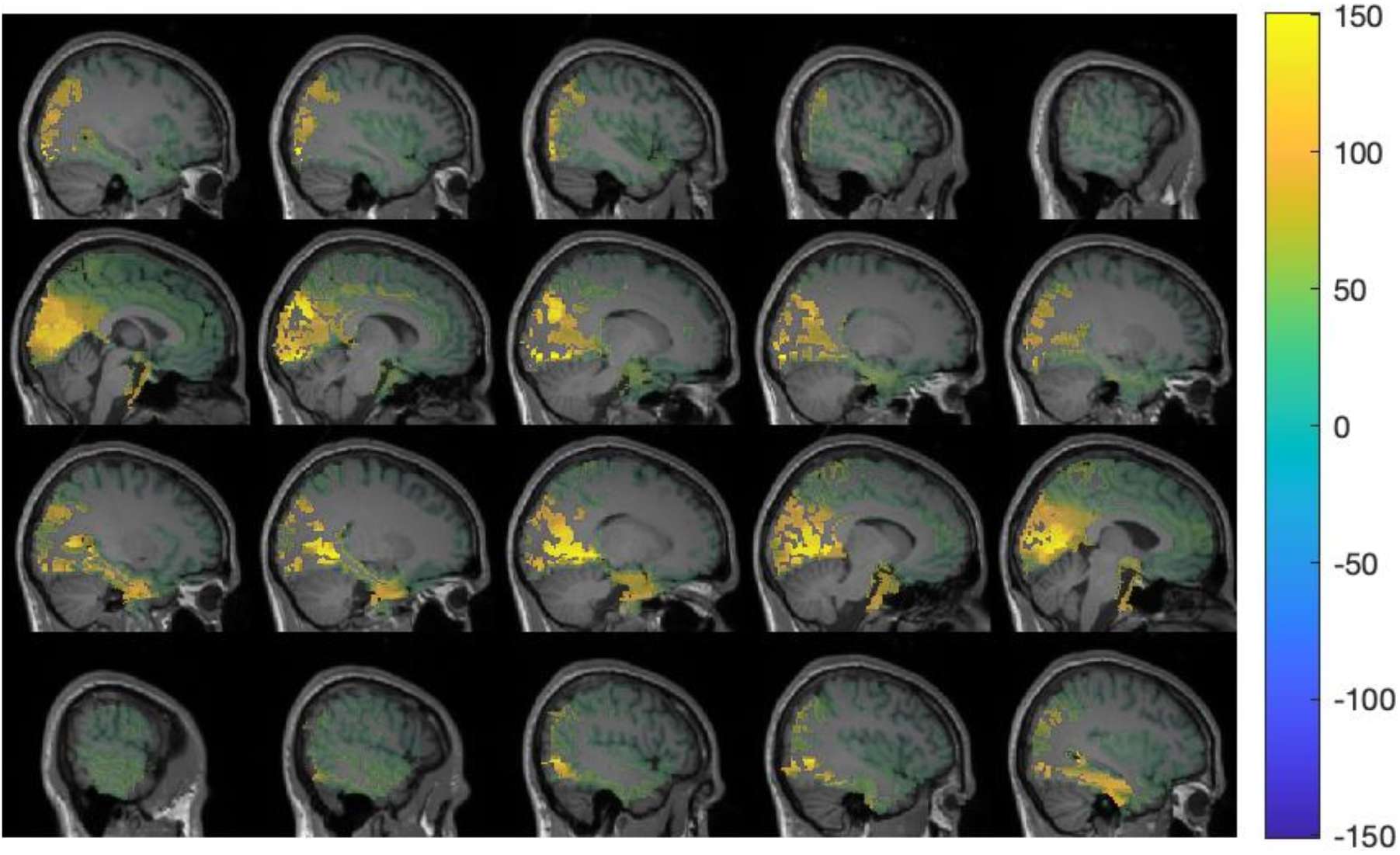
Individual-level (subject 21) contrast imaging of the alpha power during times of strong and weak occipital alpha rhythm. The colour bar refers to t-statistics indicating the mean difference between alpha power during strong and weak occipital alpha. Note this figure should not be interpreted in the context of hypothesis testing and only uses the t-statistic as a visualisation tool.

**Supplementary Figure 10.**
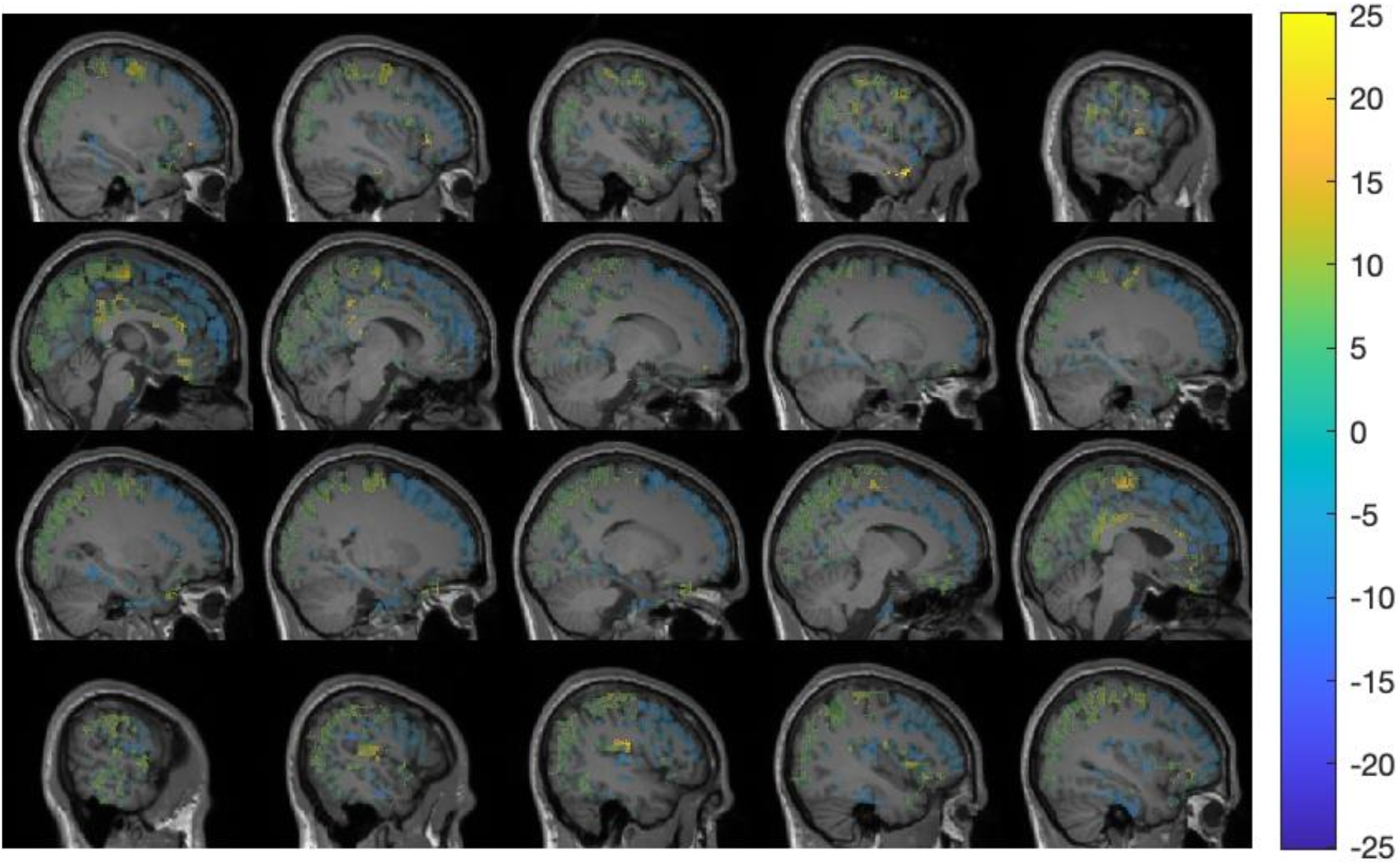
Individual-level (subject 21) contrast imaging of the excitatory to pyramidal connection strength, 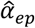, during times of strong and weak occipital alpha rhythm, with corrections for multiple comparisons applied. The colour bar refers to t-statistics indicating the mean difference between connection strength estimates during strong and weak occipital alpha.

**Supplementary Figure 11.**
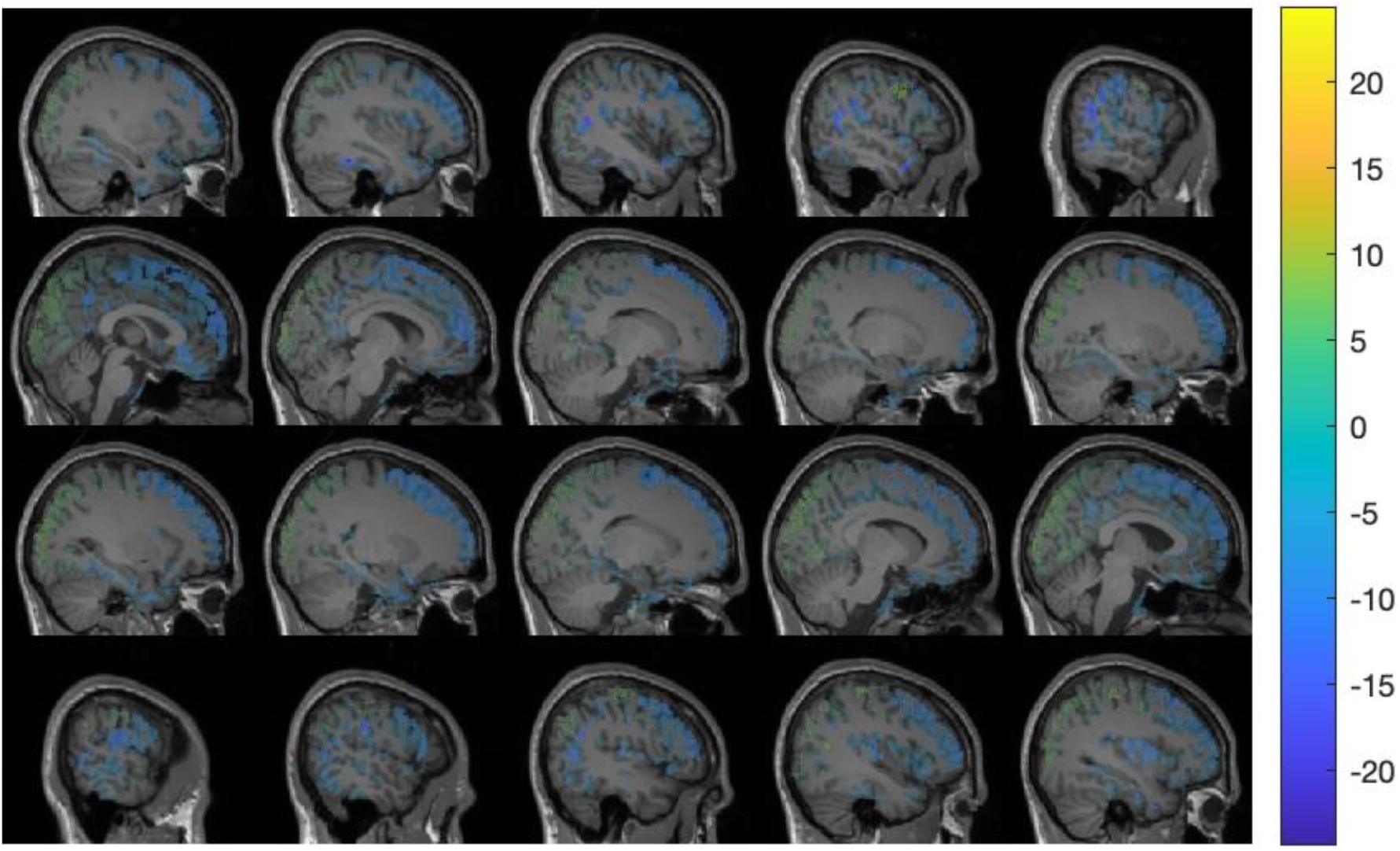
Individual-level (subject 21) contrast imaging of the inhibitory to pyramidal connection strength, 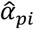, during times of strong and weak occipital alpha rhythm, with corrections for multiple comparisons applied. The colour bar refers to t-statistics indicating the mean difference between connection strength estimates during strong and weak occipital alpha.

**Supplementary Figure 12.**
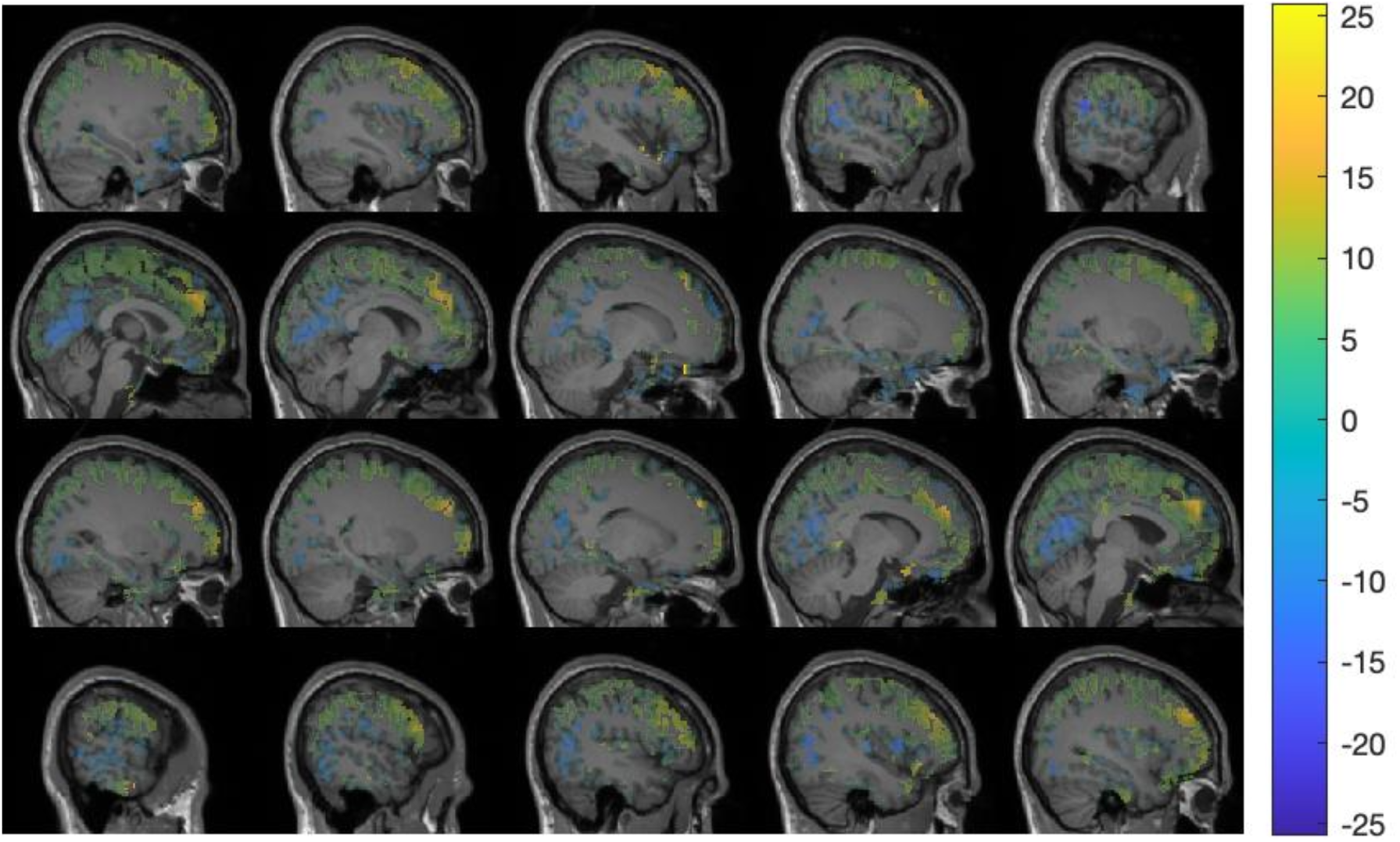
Individual-level (subject 21) contrast imaging of the pyramidal to excitatory connection strength, 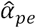, during times of strong and weak occipital alpha rhythm, with corrections for multiple comparisons applied. The colour bar refers to t-statistics indicating the mean difference between connection strength estimates during strong and weak occipital alpha.

**Supplementary Figure 13.**
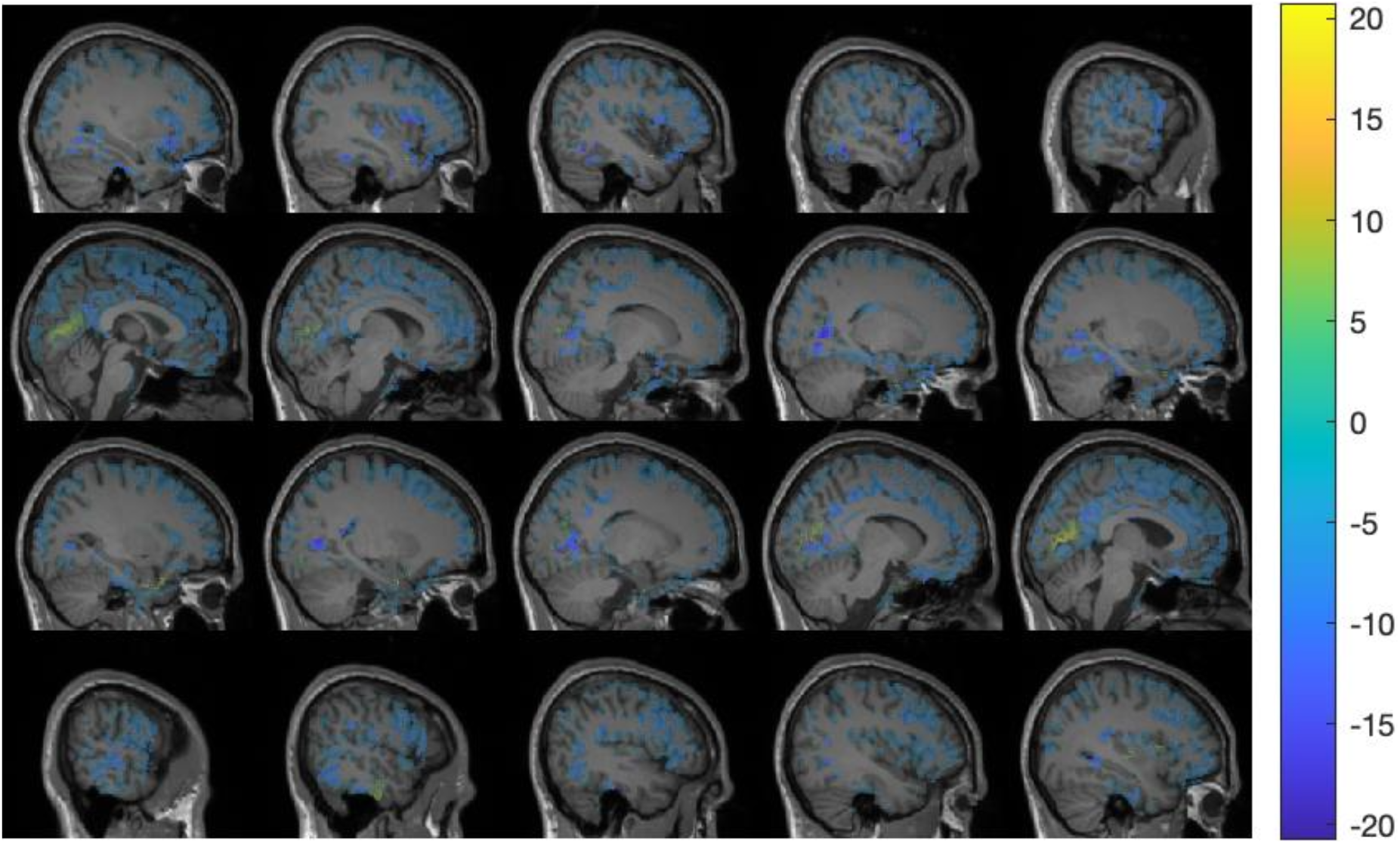
Individual-level (subject 21) contrast imaging of the pyramidal to inhibitory connection strength, 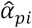, during times of strong and weak occipital alpha rhythm, with corrections for multiple comparisons applied. The colour bar refers to t-statistics indicating the mean difference between connection strength estimates during strong and weak occipital alpha.

**Supplementary Figure 14.**
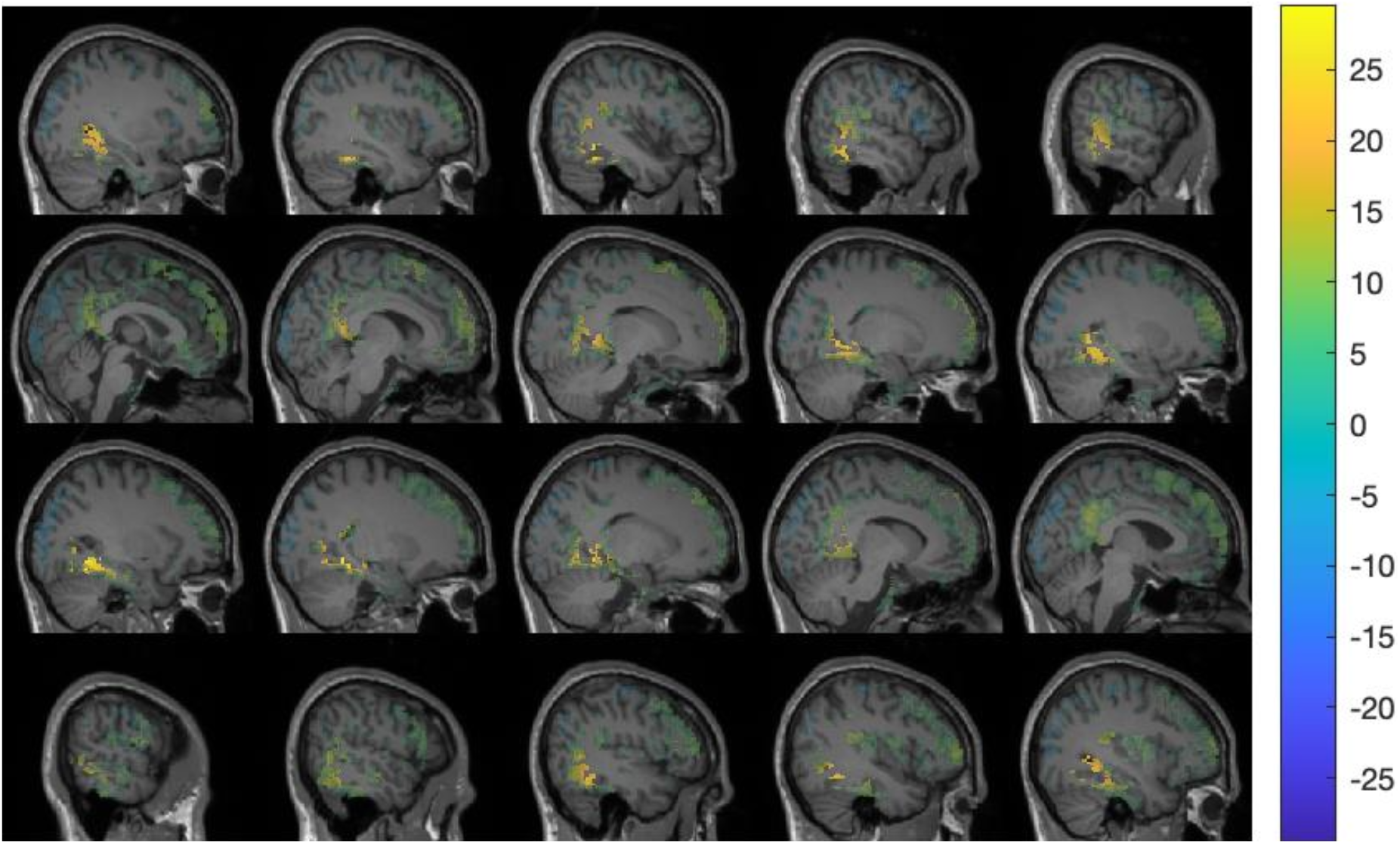
Individual-level (subject 21) contrast imaging of the external cortical input, 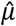, during times of strong and weak occipital alpha rhythm, with corrections for multiple comparisons applied. The colour bar refers to t-statistics indicating the mean difference between cortical input estimates under strong and weak posterior alpha.

**Supplementary Figure 15.**
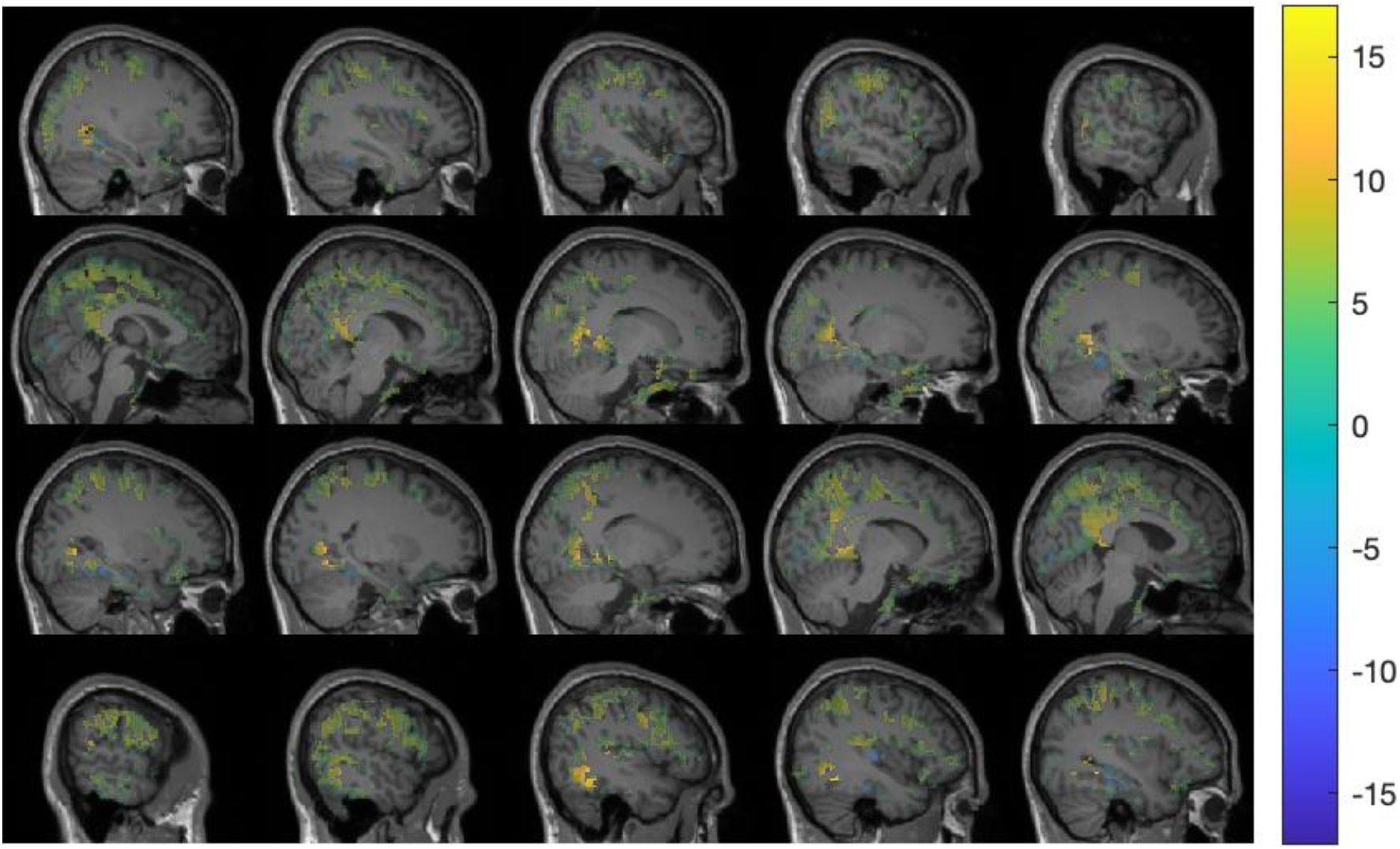
Individual-level (subject 21) contrast imaging of the membrane potential of excitatory population, 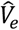, during times of strong and weak occipital alpha rhythm, with corrections for multiple comparisons applied. The colour bar refers to t-statistics indicating the mean difference between membrane potential estimates under strong and weak posterior alpha.

**Supplementary Figure 16.**
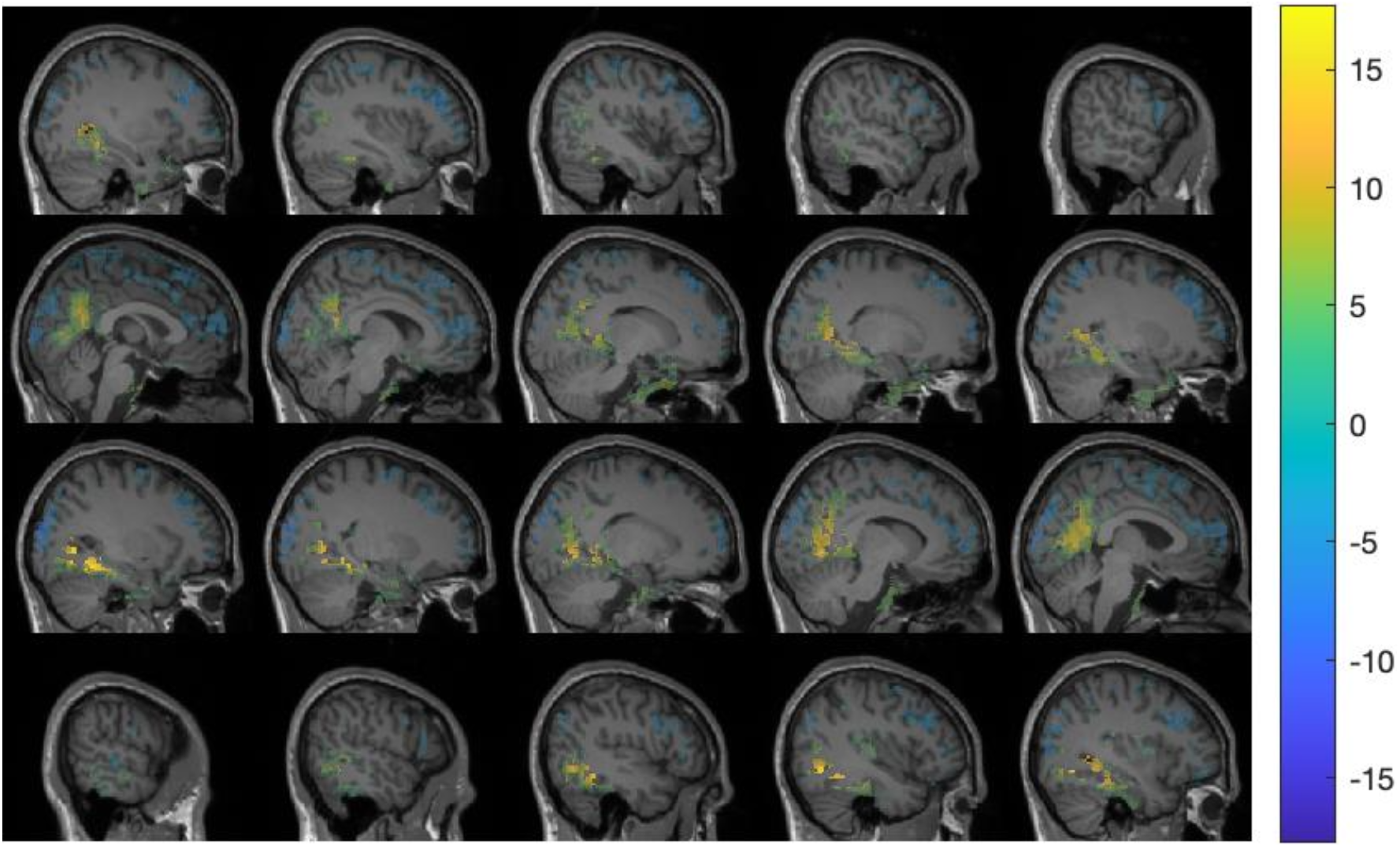
Individual-level (subject 21) contrast imaging of the membrane potential of inhibitory population, 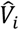, during times of strong and weak occipital alpha rhythm, with corrections for multiple comparisons applied. The colour bar refers to t-statistics indicating the mean difference between membrane potential estimates under strong and weak posterior alpha.

**Supplementary Figure 17.**
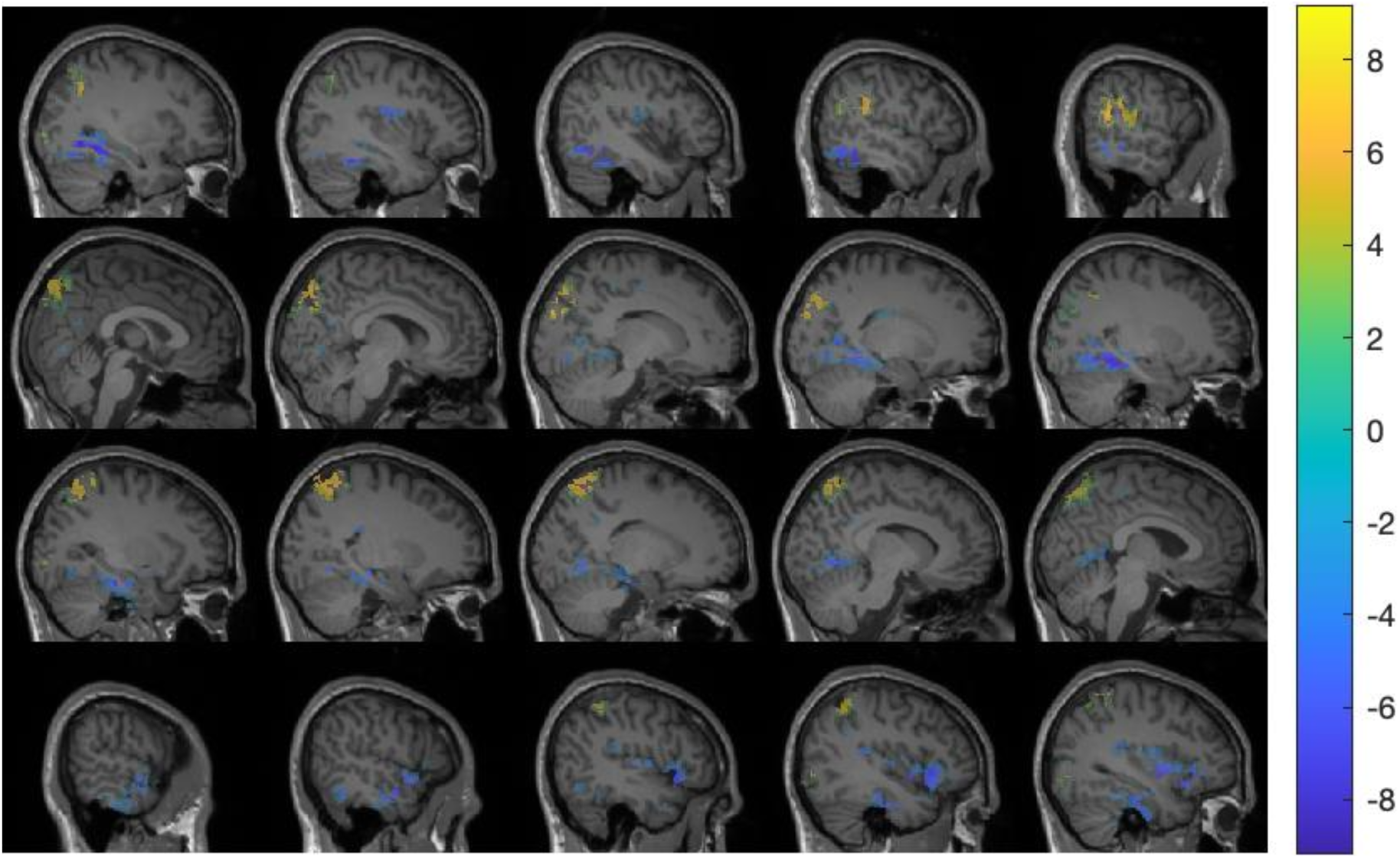
Individual-level (subject 21) contrast imaging of the membrane potential of pyramidal population, 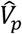, during times of strong and weak occipital alpha rhythm, with corrections for multiple comparisons applied. The colour bar refers to t-statistics indicating the mean difference between membrane potential estimates under strong and weak posterior alpha.

**Supplementary Table 1.** Top six brain regions with group-level t-statistic and coordinates obtained from each type of imaging. The group-level t-statistic with the largest absolute value was chosen from each region and presented. Region names are AAL atlas brain structure names. Only neurophysiological variables having significant group-level t-statistic source points are presented.

| Contrast imaging |  |  |  | Correlation imaging |  |  |  |
| --- | --- | --- | --- | --- | --- | --- | --- |
| Variable | Region name | t-statistic | Voxel Coordinate | Variable | Region name | t-statistic | Voxel Coordinate |
| $\mu$ | Temporal_Mid_R | 10.121 | 60,-12,-18 | $\alpha_{ep}$ | Frontal_Med_Orb_R | 9.7021 | 0,24,-12 |
|  | Calcarine_L | 9.6948 | -6,-54,6 |  | Rectus_L | 9.5168 | 0,24,-18 |
|  | Lingual_R | 9.5434 | 6,-60,6 |  | Olfactory_L | 9.3654 | -6,18,-12 |
|  | Lingual_L | 9.3446 | -12,-66,0 |  | Rectus_R | 8.6942 | 6,24,-24 |
|  | Calcarine_R | 9.3082 | 6,-60,12 |  | Frontal_Sup_Orb_L | 8.2108 | -12,18,-18 |
|  | Postcentral_L | 9.1321 | -54,-6,24 |  | Temporal_Inf_L | 7.4308 | -42,0,-42 |
| $V_i$ | Lingual_L | 18.797 | -12,-60,0 | $\mu$ | Calcarine_L | 10.4861 | -6,-54,6 |
|  | Precuneus_L | 16.0413 | -24,-54,6 |  | Calcarine_R | 9.0885 | 12,-66,6 |
|  | Precuneus_R | 14.3723 | 12,-54,24 |  | Rectus_L | -8.6522 | -6,42,-24 |
|  | Calcarine_L | 14.1709 | -24,-60,6 |  | Lingual_L | 8.3574 | -18,-66,-12 |
|  | Cuneus_L | 13.521 | -18,-60,24 |  | Frontal_Sup_Orb_L | -8.3001 | -12,42,-24 |
|  | Calcarine_R | 13.3257 | 12,-60,18 |  | ParaHippocampal_L | 8.2792 | -24,-42,-6 |
| $V_e$ | Lingual_L | 7.7466 | -18,-48,-6 | $V_i$ | Lingual_L | 16.6468 | -12,-66,-6 |
|  | ParaHippocampal_L | 6.7427 | -18,-24,-24 |  | Occipital_Inf_L | 14.2884 | -48,-60,-18 |
|  | Fusiform_R | 6.7263 | 30,-6,-36 |  | Calcarine_L | 14.1221 | 0,-66,12 |
|  | Precuneus_L | 6.6357 | 0,-54,18 |  | Cuneus_L | 14.0823 | -18,-60,24 |
|  | Olfactory_L | 6.1617 | -6,18,-12 |  | Temporal_Inf_L | 13.16 | -60,-36,-18 |
|  | Lingual_R | 6.0356 | 12,-30,-6 |  | Temporal_Inf_R | 12.9342 | 60,-42,-12 |
| $V_p$ | Lingual_R | -9.0445 | 18,-72,-6 | $V_e$ | Frontal_Inf_Orb_L | 17.7731 | -18,12,-24 |
|  | Fusiform_R | -7.5169 | 24,-72,-6 |  | Frontal_Med_Orb_R | 15.9354 | 0,24,-12 |
|  | Occipital_Inf_R | -7.3472 | 36,-78,-12 |  | Rectus_L | 15.7826 | 0,24,-24 |
|  | Calcarine_R | -7.1958 | 18,-66,6 |  | Temporal_Inf_R | 15.6033 | 48,0,-42 |
|  | Temporal_Inf_R | -6.682 | 42,-60,-6 |  | Rectus_R | 15.4561 | 6,24,-24 |
|  | Temporal_Mid_R | -6.4322 | 42,-66,0 |  | Insula_R | 15.0437 | 42,-12,0 |
| | | | | $V_p$ | Frontal_Med_Orb_R | 9.1021 | 12,42,-12 |
|  |  |  |  |  | Lingual_R | -8.2377 | 24,-48,-6 |
|  |  |  |  |  | Calcarine_R | -7.9877 | 18,-66,6 |
|  |  |  |  |  | Lingual_L | -7.8028 | -18,-72,-12 |
|  |  |  |  |  | Frontal_Inf_Orb_R | 7.4733 | 24,24,-24 |
|  |  |  |  |  | Calcarine_L | -6.9018 | -12,-60,12 |

**Supplementary Table 2.** Parameter space for filters performance exploration. Two model parameters are fixed to the model default parameter values for results visualisation. External input ranges from 640 to 820 with 20 as an interval. The inhibitory to pyramidal connection strength ranges from 640 to 820 with 20 as an interval. The excitatory to pyramidal connection strength ranges from 820 to 1000 with 20 as an interval. The pyramidal to inhibitory connection strength and the pyramidal to excitatory connection strength are fixed to 210 and 1050, respectively.

| Parameter | Lower bound | Upper bound |
| --- | --- | --- |
| $\alpha_{ip}$ | -1030 | 850 |
| $\alpha_{pi}$ | 210 | 210 |
| $\alpha_{pe}$ | 1050 | 1050 |
| $\alpha_{ep}$ | 820 | 1000 |
| $\mu$ | 640 | 820 |

